# Optogenetic WNT signaling drives germ layer self-organization in a human gastruloid model

**DOI:** 10.64898/2026.02.23.707602

**Authors:** Hunter J. Johnson, David M. McMullin, Joshua A. Zimmermann, Chang N. Kim, Nicole A. Repina, Ritu Bhalerao, Tomasz J. Nowakowski, David V. Schaffer

## Abstract

*In vitro* stem cell models of human gastrulation have been an advance for developmental biology, though elucidating mechanisms of germ layer formation remains challenging. While investigating whether spatially-patterned signaling is required for germ layer formation, we tested a “salt-and-pepper” signaling strategy in which WNT was optogenetically activated in a subset of human pluripotent stem cells (hPSC) uniformly mixed into an aggregate. Following mesendodermal specification, WNT-activated cells spatially segregated into a hemisphere, then underwent further differentiation and organization into mesoderm and endoderm. RNAseq-based lineage analysis revealed that WNT activation non-autonomously induced TGFβ/BMP signaling, leading to robust emergence of an anterior visceral endoderm-like population that patterned adjacent neural and mesendodermal fates. Transcriptional profiles and trajectories closely mirrored those observed during human gastrulation. Moreover, TGFβ or cadherin perturbation disrupted germ layer formation or spatial organization, respectively. This simple model thus enables mechanistic dissection of complex human lineage specifications and organization during gastrulation.

## Main

Gastrulation is a highly regulated and incompletely understood phase of mammalian embryo development. During this process, the initial anterior-posterior (AP) axis of the organism is established, followed by the axial arrangement of the three germ layers, i.e. ectoderm, mesoderm, and endoderm^1,2^. In mouse, AP polarity is initiated before and refined during gastrulation wherein signaling centers such as the anterior visceral endoderm (AVE) secrete inhibitors (e.g. *Dkk1, Cer1, Lefty1*) that orchestrate opposing gradients from posterior extraembryonic/epiblast domains that secrete inductive WNT, NODAL, and BMP cues^3,4^. While key biochemical and molecular factors necessary for axial establishment are thus well known *in vivo* in mouse,^4^ the spatiotemporal mechanisms that coordinate symmetry breaking and germ layer organization are unknown, especially in human.

Several hypotheses have been proposed for how morphogen signaling establishes patterns during development. One proposes that an externally imposed morphogen gradient, established by a signaling center and sink, establishes positional information based on cellular sensing of concentration thresholds^5–7^. For example, in development, graded morphogen distributions such as rostrocaudal WNT/FGF/Retinoic Acid gradients are translated into ordered neural tube patterning^8^. Analogous gradients may also be formed in gastruloid models, either by surface to interior diffusion of exogenous agonists or through engineered systems such as microfluidics or micropatterning^9,10^. In the conceptually related Turing gradient hypothesis, differential diffusion of two factors produced within a structure – an activator that stimulates its own production and an inhibitor of the activator – yield signaling gradients that break symmetry. This paradigm has been supported in particular with WNT signaling implicated in the formation of follicles, digits, and fingerprint patterns^11–13^.

In a contrasting model, “salt-and-pepper” distributions of signaling activity, in which spatially stochastic variation of signaling states within a group of cells biases lineage allocation, followed by cell migration and spatial segregation may underlie symmetry breaking^14–16^. This concept aligns with observations in blastocysts, where stochastic, salt-and-pepper expression of *GATA6* and *NANOG* within the epiblast apparently drives lineage segregation between primitive endoderm and epiblast^17,18^.

Human pluripotent stem cell (hPSC) models of gastrulation offer the opportunity to investigate such mechanisms, otherwise inaccessible in human embryos, that may regulate gastrula symmetry breaking and cell organization^19^. Recent studies have offered striking examples of hPSCs recapitulating developmental processes *in vitro* to generate patterned structures that emulate aspects of gastrulation^10,20–26^. Many of these protocols involve high concentration WNT agonist treatment in extracellular matrix materials (ECM) such as Matrigel and potentially uniform WNT activation within an organoid, after which aggregates spontaneously break symmetry and elongate into structures containing ectoderm, mesoderm, and endoderm^27–30^. However, some studies indicate that an external to internal gradient of WNT agonist or other externally added cues (e.g. ECM) could lead to polarized or radially patterned signal activation, raising questions of whether such systems encompass salt-and-pepper patterning^18,31–37^. Symmetry breaking in such systems may thus result from non-uniform WNT activation or from uniform signal activation followed by downstream gradients of factors or other internal heterogeneity. Moreover, most progress has been made with mouse cells, though the resulting structures do not fully represent the three germ layers. Also, human systems involve different peri-implantation signaling dynamics of key pathways (BMP, NODAL, NOTCH)^38^.

Here, we harness our previously developed optogenetic signaling system (optoWnt)^39,40^ to rigorously simulate and test salt-and-pepper WNT signaling within an hESC aggregate. Specifically, wild-type (WT) and optoWnt-expressing cells were mixed, aggregated, and cultured in an inert 3D material. Subsequent light illumination activated signaling in heterogeneously distributed optoWnt cells within the aggregate, which then underwent mesendodermal differentiation and spatially segregated from WT cells into hemispheres. The mesendodermal hemisphere subsequently differentiated into mesodermal T/Brachyury positive (T/*BRA^+^*) and endodermal (*SOX17^+^*) domains, that further spatially segregated such that the former surrounded the latter, and in parallel, the adjacent WT hemisphere underwent ectodermal specification. This complex patterning – emerging from simple WNT activation within a subset of spatially homogeneous cells in the absence of extracellular matrix or externally imposed gradients – thus successfully recapitulated the evolutionarily-conserved internalization of endoderm within overlaying mesoderm^41,42^, in line with recent 3D reconstruction of a Carnegie 8 embryo (CS8)^43^. Additionally, single-cell RNA sequencing (scRNA-seq) revealed cell fate complexity comparable to the inner cell mass of a CS7 human embryo^44^. Single-cell mRNA velocity and cell-cell interaction analysis revealed temporal patterns in germ layer emergence and implicated roles for TGFβ signaling and cadherin switching in pattern formation that were subsequently confirmed. In sum, our study demonstrates that simple differential cell sensitivity to a spatially homogeneous signal is sufficient to drive complex symmetry breaking, spatial organization, and lineage specification into a gastruloid.

## Results

### Breaking of Spherical Symmetry in 3D Co-Culture Aggregates

We previously demonstrated that illumination of 2D cultures of mixed WT and optoWnt cells led to mesendodermal differentiation of the latter and segregation of these two populations into a random 2D mosaic pattern^40,45^, raising the intriguing question of whether some degree of spatial segregation may also occur upon salt-and-pepper WNT activation in a more biomimetic 3D system. We mixed optoWnt hESCs (which also expressed mCherry) and WT hESCs at a 1:1 ratio into 3D cellular aggregates, which were then cultured in a biologically inert biomaterial (Fig. 1b) that we have previously shown can support 3D PSC expansion or differentiation^46^. In the dark, WT and optoWnt cells maintained a randomly dispersed mixture as determined by mCherry and Hoechst imaging (Fig.1c). In striking contrast, uniform blue light illumination led to self-organization where WT and optoWnt cells segregated into distinct hemispheres (Fig. 1a). Notably, no specific directionality in the orientation of the two hemispheres was observed among different aggregates, indicating that the segregation axis was random. In contrast, aggregates cultured in Matrigel underwent radial segregation, where optoWnt cells migrated out of aggregates, away from the WT cells, and into the surrounding gel, indicating that overly strong cell interactions with the surrounding matrix overrode structure organization (Supplementary Fig. 2).

**Fig. 1.**
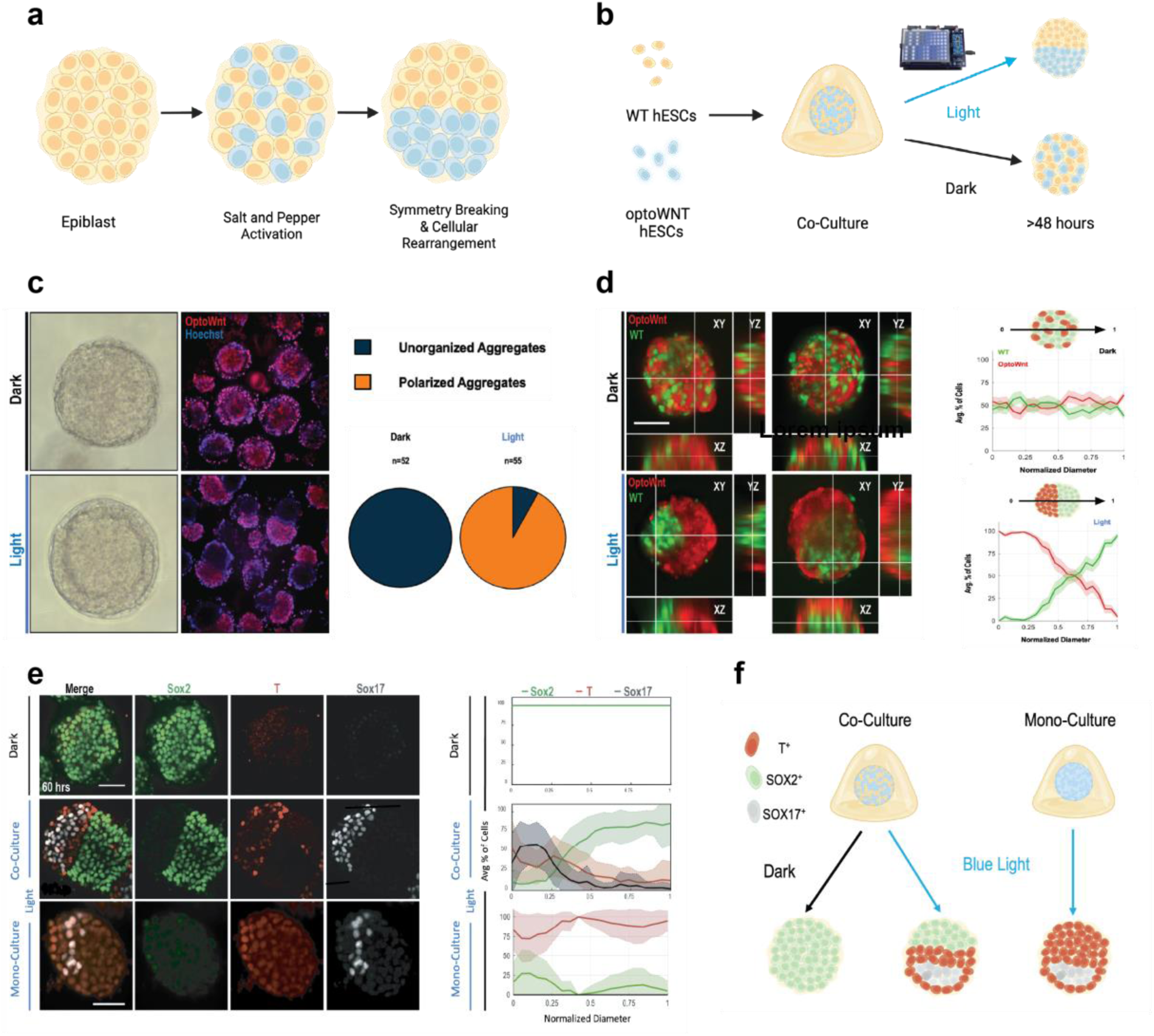
Robust segregation of WT/optoWnt hESC in 3D stem cell aggregates upon blue light stimulation. a, Schematic of the “salt-and-pepper” morphogen activation hypothesis **b,** Schematic of mixed aggregate preparation, material capture, and illumination to derive the optoWnt gastruloid model. **c,** Live cell segregation WT/optoWnt cells in 60-hour aggregates. optoWnt cells representing mCherry^+^/Hoechst^+^ and WT cells representing mCherry^-^/Hoecsht^+^. (left) Quantification of aggregate polarization, defined as deviation from mixed aggregate cell distribution in the dark (right) **d,** 3D visualization of 60-hour aggregates segregation. OptoWnt cells tagged with mCherry, WT cells tagged with GFP. (left) Line trace analysis was used to define polarized and unpolarized aggregates of optoWnt-mCherry and WT-GFP cells in the 60-hour aggregates. Scale bar = 50 μm, Error bars = S.E.M. (right) **e,** Representative maximum intensity projection images depicting germ layer patterning of 60-hour optoWnt gastruloids, as marked by Sox2 (ectoderm), T (mesoderm), and Sox17 (endoderm). (left) Line trace quantification of germ layer spatial pattering in 60-hour optoWnt gastruloids. n= 11 and 23 aggregates for dark and light respectively. (right) **f,** Model which shows the impact of co-culturing WNT activated and non-WNT wild type hESCs.

The segregation of illuminated WT and optoWnt cell aggregates was robust, as over 92% of aggregates displayed the polarized morphology (Fig. 1d), and the remaining 8% had a disproportionate number of either WT or optoWnt cells that apparently dampened self-organization. To quantify segregation, we determined the spatial vector that maximized the resolution of the WT and optoWnt cell populations for each aggregate and found that the average distribution of cell types along this vector across numerous aggregates (dark: n = 52; light: n = 55) remained mixed under dark conditions and clearly polarized under illumination (Fig. 1c). To examine cell fate, the 60-hour aggregates were stained for germ layer markers. Previously in the 2D mixed optoWnt cultures, we observed only *SOX2* and *T/BRA* expression^40^. Within the 3D aggregates, however, WT cells yielded a *SOX2*^+^ hemisphere (comprising either pluripotent or ectodermal cells), and the optoWnt region generated a core of *SOX17*^+^ endoderm within a surrounding layer of *T/BRA^+^* mesoderm (Fig. 1f). Furthermore, quantifying the distribution of *SOX2*, *T/BRA*, and *SOX17* expressing cells along the vector that maximized separation of the hemispheres demonstrated the robust segregation (>90%) of the populations under blue light conditions (Fig. 1g). This result was independent of media formulation, as segregation and germ layer marker ordering occurred in both mTeSR pluripotency medium and basal medium (Supplementary Fig.3a), though expression of pluripotency marker Nanog persisted in the WT population in hPSC maintenance but not basal media (Supplementary Fig. 3b-c). Furthermore, germ layer marker organization was found to be contingent on an optoWnt and WT cell mixture, as illuminated optoWnt mono-culture aggregates were *SOX2^-^* and the *SOX*17 subpopulation lacked organization. Finally, co-cultures kept under dark conditions retained uniform *SOX*2^+^ *T/BRA ^-^ SOX*17^-^ identity (Fig. 1e).

To investigate temporal evolution of spatial aggregate patterning, gastruloids were monitored during the 60 hours (Fig. 3a). Spheres were initially uniformly *SOX*2^+^ with no *T/BRA* or *SOX*17 expression. After 24 hours of blue light stimulation, optoWnt cells began to differentiate towards mesendoderm and express *T/BRA* yet remained well-mixed with *SOX*2^+^ WT cells, indicating that differentiation of optoWnt cells to a mesendoderm *T/BRA*^+^ phenotype precedes segregation. By 48 hours of blue light stimulation, however, distinct hemispheres of *SOX*2^+^ WT and *T/BRA*^+^ optoWnt cells had formed, but *SOX*17 was not yet present. Finally, by hour 60, the *SOX17*^+^ endodermal population emerged within the surrounding *T/BRA*^+^ mesodermal population in the mesendoderm hemisphere. RT-qPCR of bulk aggregates followed a consistent time course with *T/BRA* upregulation by hour 24, followed by downregulation of *SOX2* and upregulation of *SOX17* (Fig. 3b). Interestingly, neuroectoderm (NE) fate marker (*PAX6*) expression was also observed by hour 60, but only in illuminated conditions (Supplementary Fig. 4)^16^. Other markers of germ layer expression and epithelial to mesenchymal transition (EMT) exhibited similar temporal dynamics, with mesodermal expression peaking at 24 hours, followed by endodermal marker expression by hour 48 and maintenance at hour 60. Overall, these results support progressive temporal commitment from nascent to emergent to advanced germ layer commitment.

### Single Cell Analysis of Gastruloids Indicates Germ Layer Specification and Cell Type Complexity

Gastrulation begins when a uniform group of embryonic cells breaks symmetry through the formation of the primitive streak. This structure orchestrates differentiation of cells into the three primary germ layers (ectoderm, mesoderm, and endoderm), a critical process that serves as the foundation for all future tissues and organs^47^. To investigate the emerging cellular diversity in the optoWnt model, gene expression in 8230 cells from 60-hour gastruloids was analyzed using droplet-based scRNA-seq. As anticipated based on marker expression, uniform manifold approximation and projection (UMAP) visualization of the data revealed three general cluster areas with differential expression of canonical markers for the three germ layers (Fig. 2a and 2c).

**Fig. 2.**
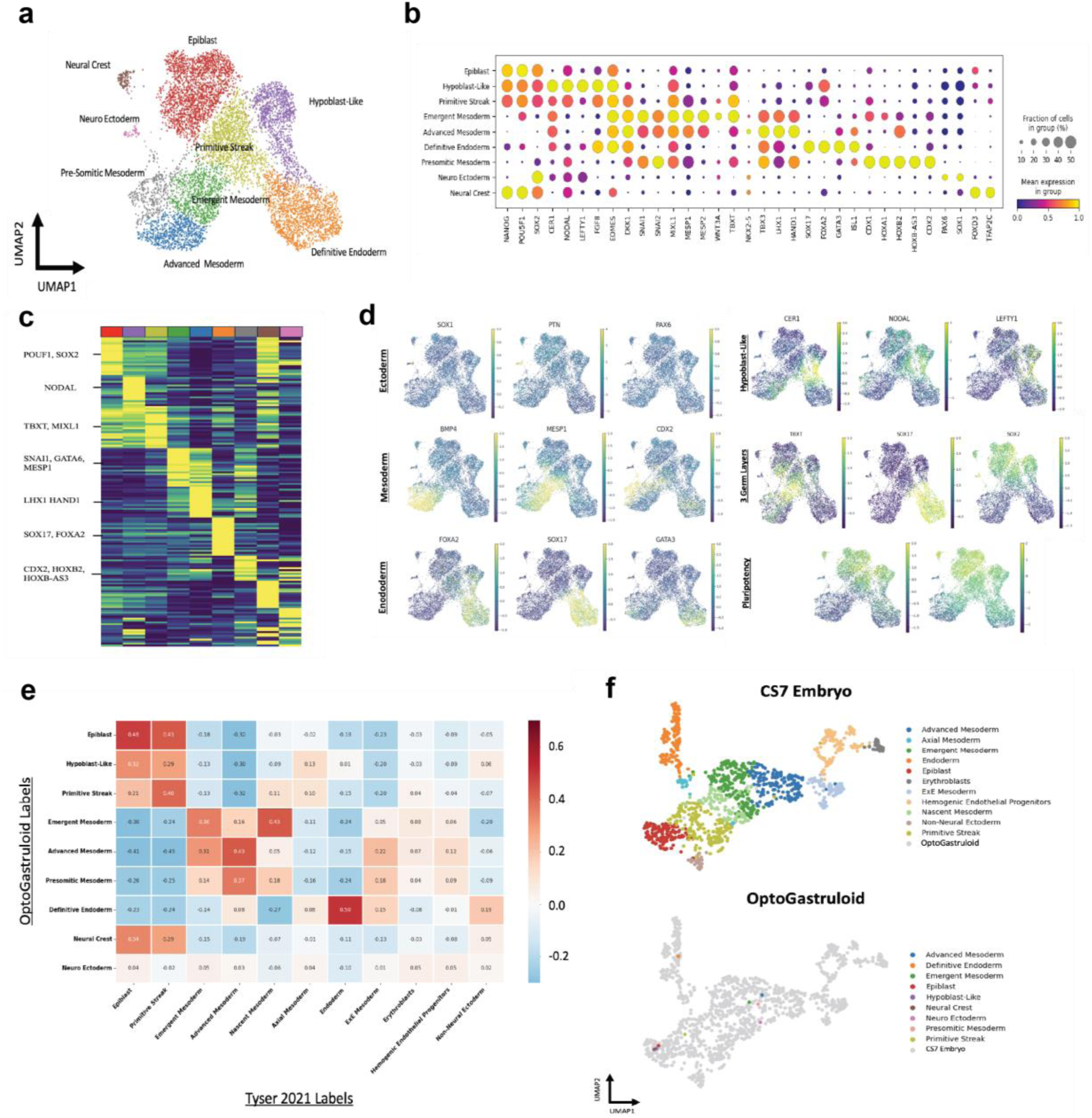
Single-cell RNA sequencing analysis of 60-hour optoWnt gastruloids with continuous blue light illumination. a, UMAP projection of scRNA-seq profiles from a pooled population of ∼50-60 hour OptoWnt gastruloids, cells are colored by unsupervised clustering. Cluster labeling was based on marker gene expression. **b,** Dot plot of mean marker gene expression level across the identified cell type specific clusters **c,** Heatmap of normalized marker gene expression; Yellow, high expression; Blue, low expression. **d,** Proportional composition of gene expression: Ectoderm (Top Left), Mesoderm (Middle Left), Endoderm (Bottom Left), Hypoblast Like (Top Right), Three Germ Layers (Middle Right), Pluripotency(Bottom Right). **e,** Cosine similarity matrix comparing OptoWnt Gastruloid Clusters (y-axis) and *Tyser 2021* cell clusters x-axis) **f,** Integration of pseudobulked OptoWnt Clusters with reference of CS7 Human Embryo^44^. See Methods. Projection of CS7 Embryo Markers (top) and projection of ∼50-60 hour optoWnt Gastruloid (bottom).

**Fig. 3.**
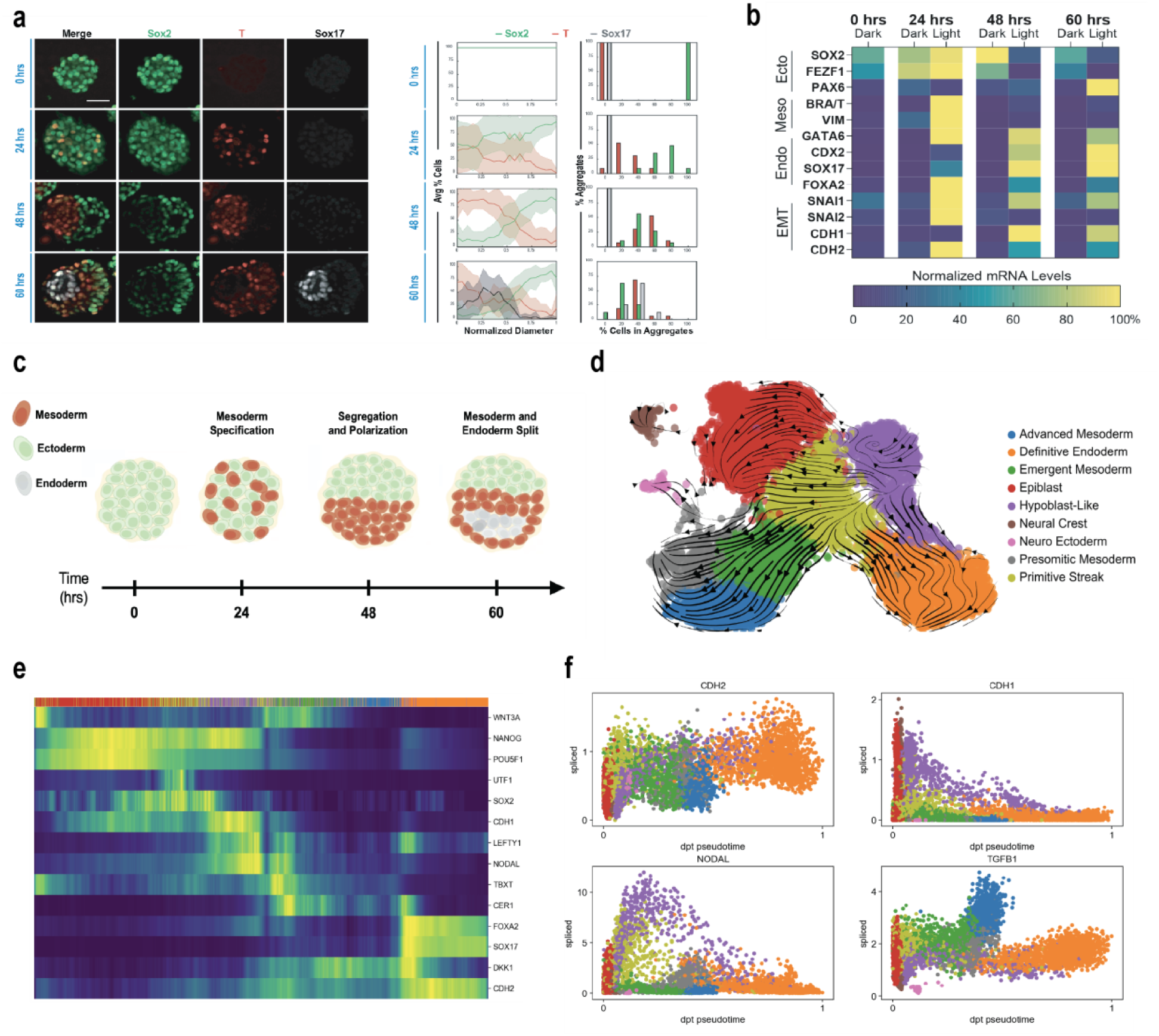
Temporal dynamics of gastruloid symmetry breaking, segregation, and germ layer emergence in the optoWnt gastruloid model. a, Representative maximum intensity projection images of temporal dynamics of germ layer emergence and aggregate polarization from 0-60 hours. (left) Line trace analysis of temporal dynamics of germ layer emergence and aggregate polarization from 0-60 hours. n= 10, 24, 31, 17 aggregates for hours 0, 24, 48, and 60, respectively. (right) (G) **b,** Heat map of RT-qPCR gene expression of germ layer markers over 0-60 hours of optoWnt gastruloid differentiation. n=3 independent preparations of ∼50 pooled aggregates. Scale bar = 50 μm. **c,** A model showing the specification and segregation of germ layers between 0 and 60 hours **d,** UMAP projection with RNA velocity vectors derived using velocyto. **e,** Heatmap of gene expression along diffusion pseudotime. **f,** Gene expression trajectories over pseudotimes. Each point represents a single cell colored by developmental state (legend in panel d).

Within each of the germ layers, we identified more specialized cell types. For instance, the ectoderm region (*SOX2^+^*) cluster harbored distinct populations of epiblast-like cells (*POU5F1*, *NANOG*, *SOX2*) and anterior visceral endoderm (AVE)-like cells (*CER1*, *NODAL*, *LEFTY1*) (Fig. 2b-d). This AVE region is a hallmark of mouse gastrulation, characterized by high expression of antagonists of NODAL, BMP, and WNT signaling (*CER1, LEFTY1, LEFTY2, DKK1*)^48,49^. This pattern also resembles the expression of the anterior hypoblast from human *in vitro* peri-gastruloids (Supplementary Fig. 5)^38^. Additionally, spontaneous generation of a neuroectoderm subpopulation of *SOX2^+^*cells is marked by the expression of *PAX6*, *SOX1*, and *PTN* (Fig. 2b-d)^50^. Furthermore, we identified a distinct subpopulation expressing markers of the neural crest (*SOX10*, *FOXD3*)^51^. These findings suggest that inhibition of NODAL and BMP by the AVE-like region may drive neural fate specification, revealing an unanticipated level of developmental complexity. We also observed a distinct *SOX17^+^* region that also highly expressed *FOXA2*, an additional canonical marker of definitive endoderm (DE)^52^. Notably, definitive endoderm populations are underrepresented in traditional high concentration WNT agonist CHIR99021-induced gastruloid models, underscoring the importance of the “salt-and-pepper” system to induce diverse signaling regimes capable of supporting complete germ layer patterning^53,54^.

The mesoderm cluster can further be segmented into distinct regions reflecting the developmental progression of various mesodermal populations. A primitive streak-like region (PS), characterized by *T/BRA* and *WNT3A* (Fig. 2b-d), marks the initiation of mesodermal differentiation and represents a transitional cell population from epiblast to nascent mesoderm and additional mesoderm lineages^55^. Also, a population of emergent mesoderm (EM) is seen in an adjacent cluster characterized by increased expression of *MESP2*, *EOMES*, and *GATA*6.

Additionally, an advanced mesoderm (AM) cluster is distinguished by high expression of *TBX3*, *NKX2-5*, *HAND1*, and *GATA6*. Notably, within the broader mesodermal population we observed clear populations expressing *CDX2^+^* and *GATA6^+^*, which correlate with distinct anterior and posterior patterning^29^. Furthermore, within the *CDX2^+^* population, there is a clear population of presomitic mesoderm (PSM) defined by expression of *TBX6*, *HES7*, and various *HOX* genes (Supplementary Fig. 4). Moreover, we observed the presence of *HES7*, a key component of the segmentation clock that regulates the rhythmic timing of somite formation^37^. We also found sparse expression of markers for dorsoventral (*PAX1*) and rostrocaudal (*UNCX*, *TBX18*) somites, indicating early stages of somite patterning. Recent work has shown that an early pulse of retinoic acid with the later addition of Matrigel can yield posterior tissues and segmented somites in a human gastruloid that closely resemble those *in vivo*^53,56^, though such lineages have not previously been observed in a simple system involving only salt-and-pepper WNT induction in cell clusters within an inert material.

Individual marker expression in the gastruloid scRNA-seq dataset was similar to expression profiles in a CS7 human gastrula^44^. Specifically, by integrating cell type labeled data from each of the CS7 gastrula and our 60-hour gastruloids, evaluation of the UMAP analysis (Supplementary Fig. 6) revealed strong alignment of endoderm, mesoderm, primitive streak, and epiblast labeled cell populations. Furthermore, we generated averaged pseudo-cells for each cell type and germ layer from our 60-hour gastruloids (Fig. 2f), which again showed good alignment with CS7 gastruloids. In sum, this scRNA-seq analysis demonstrates commitment to the three germ layers as well as biologically relevant cell type complexity in this simple optoWnt gastruloid model, underscoring its potential to recapitulate key events in early embryonic development.

### Temporal Dynamics of Gastruloid Segregation and Patterning

While WNT/β-catenin signaling is known to be critical for driving primitive streak formation, mesendoderm differentiation, and egress of these cells from the epiblast during mammalian gastrulation, the temporal sequence of these events in human gastrulation is not well understood^55,57^. We therefore analyzed the temporal dynamics of cluster commitment and cellular emergence within this model. For RNA velocity analysis, scVelo^58^ calculates velocity vectors based on quantities of spliced and unspliced transcripts, which it then projects onto the annotated UMAP (Fig. 3d). Velocity streamlines originated from two primary sources: epiblast cells and the AVE-like region. From the epiblast, vectors primarily progressed toward an early primitive streak population marked by *T*/*BRA^+^/SOX2^+^* and *MESP1⁻,* which subsequently flowed toward neuroectoderm^59^. Vectors originating in the AVE-like region bifurcated, extending into the primitive streak and continuing to the presomitic mesoderm as well as to the definitive endoderm. An intermediate mesendodermal state of the primitive streak co-expressing *NODAL*, *EOMES*, and *T/BRA* displayed two main vectors (Supplementary Fig. 8). One population extended towards emergent and Advanced Mesoderm, and a second through the AVE-like cells and the DE. Similar transitional states have been described *in vivo* where *EOMES* expressing posterior epiblast cells in the mouse embryo segregate into mesoderm-restricted (*EOMES*/*MESP1*) and definitive endoderm restricted (*EOMES/FOXA2*) lineages, with NODAL signaling driving the balance toward the latter(Fig. 3e)^60,61^. We additionally observed vectors linking presomitic mesoderm and neuroectoderm, in agreement with a neuromesodermal progenitor (NMP) program reported to generate both spinal cord and somites in vertebrates.

While definitive *in vivo* evidence connecting presomitic mesoderm directly to neuroectoderm is lacking, similar relationships have been observed in hPSC differentiation models, where NMP-like *SOX2^+^* & *T/BRA^+^* cells generate both paraxial mesoderm and neural progenitors expressing *SOX1* and *PAX6*^37,62^.

To capture developmental ordering and cellular maturity, we employed diffusion pseudotime analysis^63^, which arranges cells along a continuous lineage path using the epiblast as a root population to reveal the sequential emergence of germ layer–specific states. Cells at early pseudotime were characterized with canonical genes of pluripotency *OCT4*, *POU5F1, SOX2*, and *NANOG.* This was followed by the induction of an EMT marked by *SNAI1/SNAI2* as well as a cadherin switch from *CDH1* to *CDH2* alongside transitional markers *T/BRA*, *NODAL*, and *WNT3A* (Fig. 3f and Supplementary Fig. 8). From this transitional state, later pseudotime showed strong expression for more mature cell types including *SOX17* and *FOXA2* for definitive endoderm, *MESP2* and *TBX6* for presomitic mesoderm, and *PAX6* and *SOX1* for neuroectoderm (Fig. 3e). These observations align with previous studies and recapitulate the ordering of key features of human gastrulation including EMT-like transition and germ layer emergence and maturation^64^. Together, the pseudotime and RNA velocity-based analysis depict clear developmental trajectories that closely align with prior studies of human development.

### Germ Layer Segregation and Endoderm Differentiation is Dependent on TGFβ, ACTIVIN, and NODAL Signaling

*In vivo* and *in vitro*, the TGFβ superfamily and WNT cooperate to differentiate pluripotent cells into the primitive streak and subsequently the mesendoderm^18,65–69^. WNT activity induces and sustains expression of TGFβ1-3, *ACTIVIN*, and *NODAL*, which in turn activate SMAD2/3. The duration and magnitude of this SMAD2/3 activation bias subsequent fate decisions, where sustained signaling promotes definitive endoderm (DE) and transient signaling supports mesoderm^70,71^. To identify candidate signaling patterns that may influence gastrulation, we performed ligand receptor inference using CellChat, a computational tool developed to infer intercellular communication within scRNA-seq datasets (Supplementary Fig. 9-11)^72^. The predicted strength of outgoing and incoming ligand-receptor interaction strengths between the labeled cell types revealed distinct TGFβ superfamily signaling activities among different subpopulations (Fig. 4a-b). Specifically, Emergent Mesoderm and Advanced Mesoderm apparently strongly send ACTIVIN signaling that is received by epiblast, the AVE-like region, and the Definitive Endoderm populations. Conversely, NODAL is predominantly sent by the Epiblast and AVE-like region and received by Advanced Mesoderm, Emergent Mesoderm, and Definitive Endoderm. Moreover, TGFβ-1 signaling is strongly sent from the advanced mesoderm and strongly received by the AVE-like region and the Definitive Endoderm. Notably, among the TGFβ superfamily ligands signaling through ALK4/5/7, only NODAL is predicted to be sent by cell populations present at the onset of differentiation (Epiblast and AVE-like cells), whereas TGFβ-1 and ACTIVIN are predominantly sent by mesodermal populations that emerge later. This suggests that epiblast-derived NODAL is the primary ALK4/5/7 ligand driving initial mesendoderm specification, whereas TGFβ and ACTIVIN signaling from mesodermal populations subsequently contributes to endoderm differentiation and segregation.

**Figure 4:**
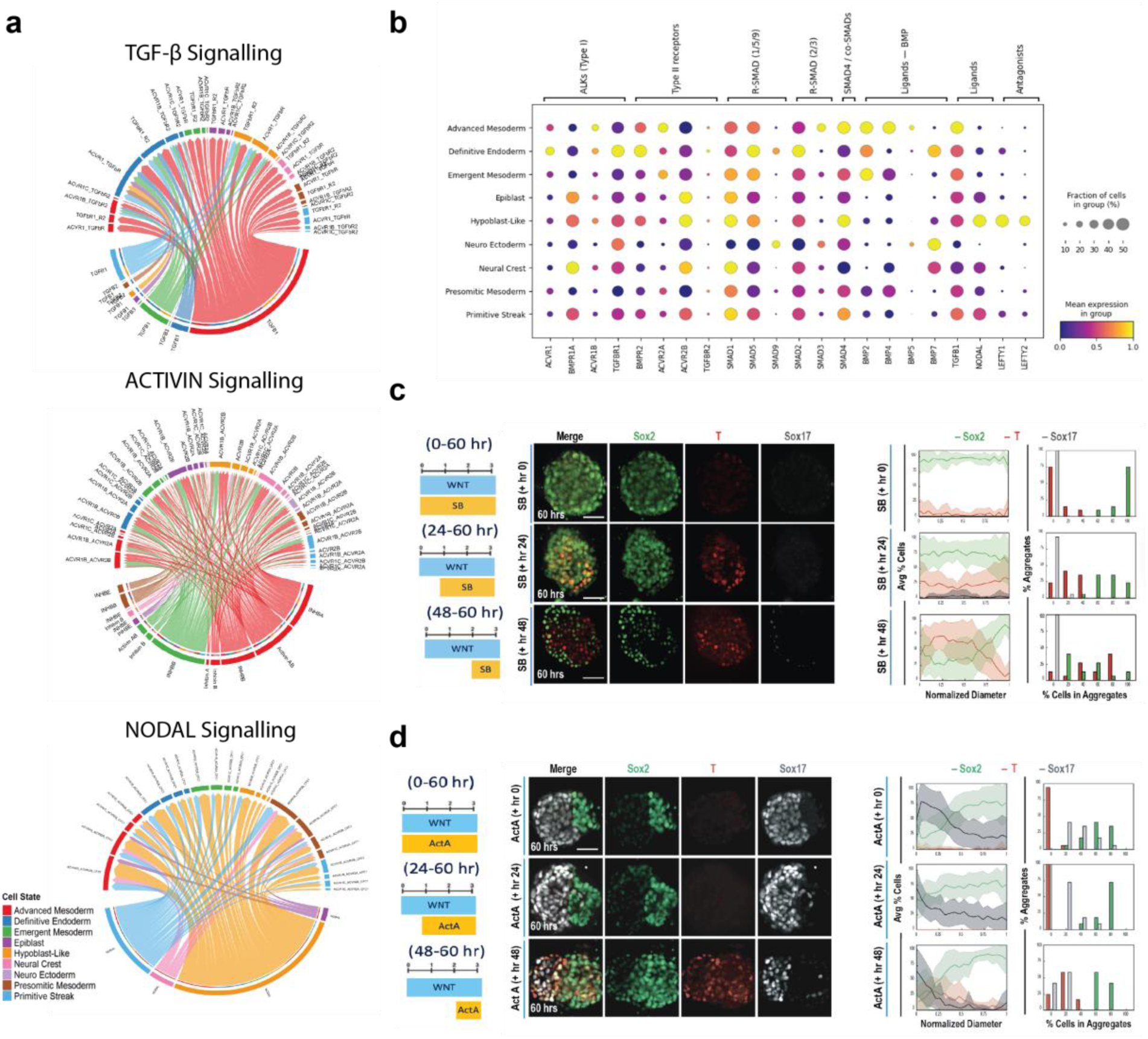
optoWnt gastruloid germ layer emergence is dependent on transient TGFβ signaling in the optoWnt population. **a,** Chord diagram depicting Ligand – Receptor (L-R) for TGfβ (Top), ACTIVIN (Middle), and NODAL(Bottom) derived from CellChat pathway analysis. **b,** Dot plot of mean marker gene expression level for TGfβ pathway related genes. **c,** Representative maximum intensity projection images of germ layer emergence with TGFβ pathway inhibition via addition of SB431542 (10 μM) at either hour 0, 24, or 48 of 60-hour gastruloid light illumination. (left) Quantification of germ layer patterning and population distribution with TGFβ pathway inhibition via addition of SB431542 (10 μM) at either hour 0, 24, or 48 of 60-hour gastruloid light illumination. n= 21, 18, 16 gastruloids for SB addition at hour 0, 24, and 48 respectively. (right) **d,** Representative maximum intensity projection images of TGFβ pathway activation via addition of Activin-A (50 ng/mL) at either hour 0, 24, or 48 of 60-hour gastruloid light illumination. (left) Quantification of germ layer patterning and population distribution with TGFβ activation via addition of Activin-A (50 ng/mL) at either hour 0, 24, or 48 of 60-hour gastruloid light illumination. n= 18, 11, 13 gastruloids for Activin-A addition at hour 0, 24, and 48 respectively. Scale bars = 50 μM. Error bars = std. dev. (I) Model of TGFβ activation and germ layer specification. (right)

This signaling pattern also aligned well with expression of other TGFβ superfamily members from the scRNA-seq dataset To test some implications of these computational predictions, TGFβ signaling through ALK4/5/7 was inhibited using the small molecule SB-431542, which blocks signaling from all three TGFβ isoforms and *NODAL*. Inhibitor treatments started at 0, 24, or 48 hours, and cells were fixed and stained at 60 hours (Fig. 4c left image). Treatment from 0-60 hours blocked mesendoderm differentiation of optoWnt cells, resulting in *SOX2^+^ SOX17^-^* aggregates with very minimal *T/BRA* expression (Fig. 4c top row), illustrating that initial mesendoderm differentiation depends on WNT-mediated TGFβ signaling. Interestingly, inhibition starting at 24 hours permitted the emergence of *T/BRA^+^* mesendoderm optoWnt cells yet blocked subsequent cell segregation and *SOX17* expression (Fig. 4c middle row). Furthermore, inhibition at 48 hours impaired but did not fully eliminate *SOX17^+^* cell differentiation, yet limited polarization of *T/BRA* and *SOX2* (Fig. 4c, bottom row). Together, these results demonstrated the necessity of TGFβ signaling not only for initial mesendoderm differentiation but also for segregation of WT and optoWnt cells, a result consistent with the known role of TGFβ in EMT^73^. Interestingly, a subpopulation of optoWnt cells committed towards endoderm in a TGFβ-dependent manner during the cell sorting timeframe, since inhibition of TGFβ at hour 24, but not hour 48, inhibited the emergence of a *SOX17*^+^ population at 60 hours.

We also activated TGFβ signaling via the addition of Activin A at 0, 24, or 48 hours of the 60-hour time course. In contrast to inhibition of TGFβ with SB-431542, Activin A addition at 0 and 24 hours still enabled hemisphere segregation, yet resulted in uniform *SOX17^+^* and *T/BRA^-^*expression within the optoWnt hemisphere (Fig. 4d top & middle row), consistent with the combined role of WNT and TGFβ in endoderm formation^4,65^. However, addition of Activin A at 48 hours did not inhibit *T/BRA* expression at 60 hours (Fig. 4d bottom row), indicating that optoWnt cells had committed to either mesoderm or endoderm precursors by 48 hours.

WNT-driven TGFβ/ACTIVIN/NODAL signaling through ALK4/5/7 is required for initiating mesendoderm differentiation, and our gastruloid time course experiments indicate that the duration of signal exposure impacts cell fate. In the prior scRNA-seq analysis, DE populations retained high expression of TGFβ/ACTIVIN/NODAL receptors *ACVR1* (ALK2), *ACVR1B* (ALK4), *TGFBR1* (ALK5), *TGFBR2* alongside *SMAD2,* a pattern that likely supports sustained SMAD2/3 activation by TGFβ/ACTIVIN/NODAL (Fig. 4b)^74,75^. In contrast, mesodermal populations displayed elevated *ACVR1B* (ALK4), *ACVR2A*, and *SMAD3* but reduced *TGFBR1*, *TGFBR2*, and *SMAD2*, indicating a potential shift toward SMAD1/5/9 signaling marked by increased expression of *SMAD1/5* and *BMP2/4* ^76^. This shift could arise from receptor competition, as ACVR2A can complex with ALK4 to mediate SMAD2/3 signaling or with BMP type I receptors (ALK2/3/6) to initiate SMAD1/5/9 signaling when BMP ligands are present^70^. This interpretation is reflected in our perturbations where ALK4/5/7 inhibition at 0 hours blocked all mesendoderm, at 24 hours prevented DE but allowed mesoderm commitment, and at 48 hours yielded very sparse DE. Conversely, Activin A addition biased differentiation toward DE and suppressed *T/BRA*. These dynamics parallel live-cell studies showing that transient SMAD2 nuclear translocation is “remembered,” and prior WNT exposure alters transcriptional competence^71^. Together, these results implicate the temporal role of TGFβ/ACTIVIN/NODAL signaling dynamics and germ layer specification in the optoWnt gastruloids, where continuous TGFβ activation instructs the endoderm and transient TGFβ activation instructs the mesoderm.

### Spatial Segregation of Germ Layers is Mediated by Differential Cadherin Expression

Throughout the optoWnt gastruloid development process, cells migrate, segregate, and self-organize into regions in coordination with germ layer differentiation. To study mechanisms underlying this intrinsic cellular organization and dynamic polarization, we investigated cell adhesion pathways. Differential adhesion mediated through expression of various cell-cell adhesion proteins (e.g. cadherins) has long been known as a mediator of cell sorting phenomena^67,77^. Furthermore, cadherin expression is dynamically regulated during gastrulation and has been recently used to enhance cell sorting of synthetic embryos^78^. Epiblast and pluripotent cells express E-cadherin (*CDH1*), but upon differentiation into the primitive streak, they undergo an epithelial-to-mesenchymal transition, downregulate E-cadherin, and initiate N-cadherin (*CDH2*)^79–81^. Here, we likewise observed differential cadherin expression in the 60-hour gastruloids (Fig 5e and Supplementary Fig 12), with *CDH1* expression in the epiblast and AVE-like populations and *CDH2* expression highest in the mesoderm and endoderm population (Fig. 5d). Furthermore, optoWnt cells under blue light stimulation upregulated *CDH2* expression by 24 hours and lost *CDH1* expression by 48 hours (Fig. 5 a-b), suggesting that cadherin-mediated sorting could play a role in the polarization of co-culture aggregates. Time course evaluation revealed that populations of *CDH1^+^* and *CDH2^+^*cells progressively polarized, initially observed at 24 hours, with robust segregation into two clear hemispheres by hour 60.

**Fig. 5.**
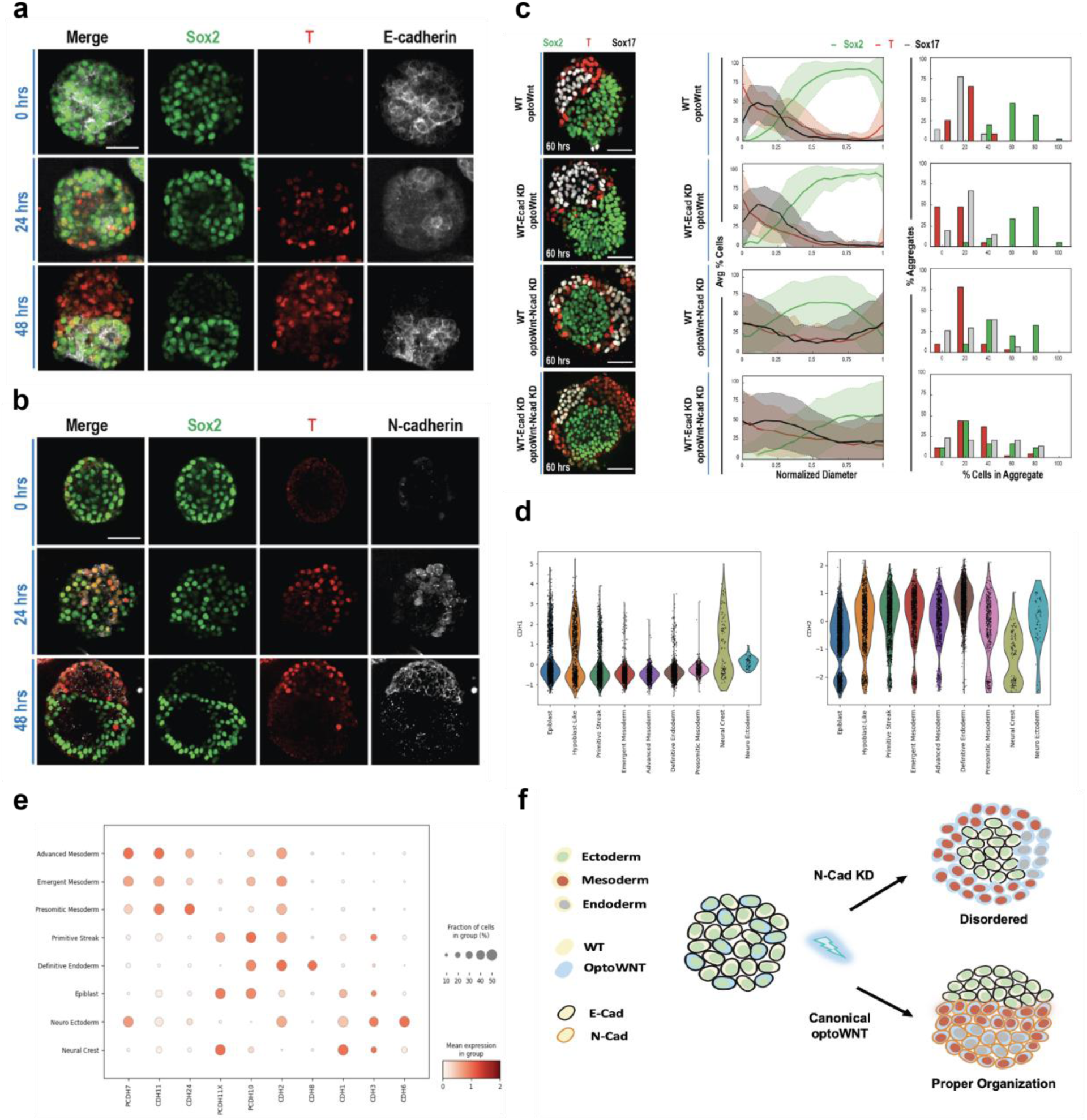
optoWnt gastruloid segregation is dependent on cadherin switching and an EMT-like event. a, Representative maximum intensity projection images of germ layer temporal dynamics and E-cad expression in the optoWnt gastruloid pre and post polarization (0-48 hours). **b**, Representative images of germ layer temporal dynamics and N-cad expression in the optoWnt gastruloid pre and post polarization (0-48 hours). **c,** Representative maximum intensity projection images of germ layer patterning in the 60-hour optoWnt gastruloids with varying mixtures of WT and optoWnt hESC knockdown lines. (left) Quantification of germ layer spatial patterning and population distribution in the 60-hour optoWnt gastruloids with varying mixtures of WT and optoWnt hESC knockdown lines. n= 60, 22, 45, 44 gastruloids for each respective condition (top-down). Scale bars = 50 μM. Error bars = std. dev. (right) **d,** Expression of E-Cadherin (CDH1) and N-Cadherin (CDH2) within cell type clusters **e,** Dot plot of expression levels of various cell adhesion genes for each cell type. **f,** Model representation of Cadherin switching and germ layer specification in both Canonical optoWNT and optoWNT N-Cad knockdown.

We tested whether *CDH1* and/or *CDH2* expression was necessary for germ layer organization within optoWnt gastruloids. Using lentivirally delivered shRNAs, *CDH1* or *CDH2* was knocked down in WT or optoWnt cells, respectively, and knockdown (KD) was confirmed at the protein level (Supplementary Fig. 12). Co-culture of optoWnt cells with WT cells lacking *CDH1* expression (WT-Ecad KDs) resulted in aggregate polarization and patterning of 60-hour gastruloids similar to the prior WT/optoWnt cultures (Figure 5c). Loss of *CDH1* expression in theWT population thus did not impair the differentiation of optoWnt cells to mesodermal or endodermal populations, establishing that cell differentiation and organization did not rely on *CDH1* alone. In contrast, in co-cultures of WT cells with optoWnt *CDH2^-^*, the latter still differentiated into mesodermal and endodermal lineages. However, the hemispherical organization was replaced with radial segregation (Fig. 5c), where a *SOX2^+^* core was surrounded by a *T/BRA^+^* and *SOX17^+^* population that failed to segregate from one another. This demonstrates that N-cadherin–mediated adhesion is required for spatial sorting of definitive endoderm from mesoderm, a mechanism supported by previous *in vitro* and *in vivo* studies showing mixed-cadherin endoderm progenitors physically sort away from mesodermal neighbors via homophilic *CDH2* interactions^52^.

Finally, a co-culture of WT *CDH1^-^* and optoWnt *CDH2^-^*cells disrupted coordinated cell segregation and random pockets of *SOX2^+^*, *T/BRA^+^*, or *SOX17^+^* populations were present throughout each aggregate without hemispherical or radial symmetry (Fig. 5f). We hypothesize that differential adhesion or lineage-specific motility within the *SOX17^+^* compartment (AVE-like and DE) could further fragment tissue organization in the absence of both E- and N-cadherin.

Other cadherins, including *CDH3* (P-cadherin) enriched in mesoderm and *CDH11* enriched in both mesodermal and endodermal lineages, may partially compensate in cell segregation as described in other developmental contexts^82,83^. In addition, E-cadherin loss may alter β-catenin availability, modifying both adhesion strength and WNT signaling, which could influence migratory behavior and morphogen interpretation^84^. Overall, these results implicate cadherin switching as critical for self-organization in human gastruloids.

## Discussion

Several models have been proposed to explain how symmetry is broken and order emerges in early development, including morphogen gradients, Turing activator-inhibitor networks, and the more recently metabolic gradient hypothesis^5,12,85,86^. *In vitro* human stem cell systems offer a key resource in testing these concepts, especially as species differences make it difficult to extrapolate directly from mouse models of gastrulation to human^38^. Here, we have explored the emergence of complexity from simplicity, i.e. the activation of WNT signaling in a subset of cells leads to the formation and self-organization of the three germ layers and development into numerous additional lineages in the absence of external extracellular matrix or other patterned cues. We propose a two-step process supported by our results. First, WNT activation in a subset of spatially well-mixed cells can partition the epiblast into *NANOG/SOX2* expressing ectoderm versus *GATA6* expressing mesendoderm, establishing the initial anterior posterior polarity of the system^22^. This framework is consistent with recent lineage-tracing work in human gastruloid-like models demonstrating that early heterogeneity in morphogen responsiveness can bias subsequent positioning and fate allocation through cell rearrangements^87^. Second, within the mesendoderm compartment, WNT and NODAL function as local activators that induce secretion of the long-range inhibitors *CER1, LEFTY*, and *DKK1*. The resulting network yields a single, stable definitive endoderm (*FOXA2, EOMES*) territory adjacent to a complementary mesodermal (*T/BRA, MESP1*) domain reminiscent of activator–inhibitor logic^18,61^.

Numerous studies have developed elegant gastruloid models based on patterning via external, spatially organized cues, including targeted manipulation of signaling centers^5,7^ and spatially heterogeneous WNT activation in hPSC gastruloids^10,32,54,88^. Others have mixed pre-differentiated cell lineages to promote self-organization, through a combination of embryonic and extraembryonic stem cells^25,89,90^, reprogramming to naïve pluripotency^24,91^, and inducible lineage-specific transcription-factor expression^23,90^. Exogenous patterned signals can thus clearly promote the development of patterned gastruloids^26–30,92,93^. However, while such gradients do clearly exist in development and are sufficient for patterning, in parallel it is also important to continue to rigorously test whether stochastic signal activation followed by cell sorting within gastruloids contributes to symmetry breaking, i.e. the extent to which spatially heterogeneous exogeneous cues are necessary for symmetry breaking. Within a spherical aggregate, the diffusion of extracellular cues into the aggregate may generate signal gradients, and ECM in general and Matrigel in particular are known to polarize numerous organoid systems (including intestinal, neural, somite, and other organoid systems), both of which may lead to spatial patterning of cell fate^33–37^. In support of this idea, in mESC gastruloids radial *Sox2* gradients emerge during the CHIR pulse with higher levels in the gastruloid core. *T/Bra* subsequently became expressed in a salt-and-pepper pattern but at higher levels in the *Sox2*^-^ periphery^94^. This observation indicates that heterogeneity in *SOX2* vs. *T/BRA* expression may arise in part from spatial gradients of cues. However, other studies have found more variable distributions of *T/BRA*, ranging from homogeneous salt-and-pepper to radially polarized, though differences in culturing conditions including the addition of ACTIVIN and FGF likely contribute to this variability^95^. In a related study, a hESC-based gastruloid model with uniform BMP4 exposure initially generated an asymmetric gradient of *SOX2* vs. *T/BRA* along an AP axis in the absence of any observable cell sorting, a process attributed to the development of internal WNT-DKK1 gradients^26^. Together, these studies establish that extracellular signaling gradients and pathway crosstalk within aggregates can generate the initial heterogeneity from which spatial patterning emerges, raising a question for the role of salt-and-pepper fate specification and self-organization. Our results, based on simple light activation of WNT (which penetrates 100s of microns into tissue) within a subset of cells within an inert biomaterial, offers a direct causal demonstration that salt-and-pepper WNT signaling alone is sufficient to drive the development and organization of the three germ layers.

Alongside these extrinsic mechanisms, accumulating evidence points to intrinsic cell to cell variability as a route to fate diversification. WNT reporters reveal a progression from uniform low activation to uniform high activation before cell-to-cell heterogeneity emerges, linked to earlier differences in NODAL and BMP signaling that modulate the duration of each cell’s WNT response^54^. Salt-and-pepper heterogeneity in fate markers arises spontaneously in naïve human ES cells^22^, and analogous stochastic variability has been observed in the inner cell mass of cultured human embryos between E5 and E7 for hypoblast and epiblast cells^96^. Consistent with this intrinsic variability, endoderm emerges from *T/BRA*^+^ mesendoderm with a salt-and-pepper distribution during mESC gastruloid development^97^. Similar principles of cell-to-cell heterogeneity extend beyond early development into for example the activity of YAP1 as a function of tissue density^98^. Collectively, these observations demonstrate that intrinsic stochastic fluctuations in morphogen sensitivity and stem cell state can contribute alongside or conceivably instead of extrinsic spatial cues in generating fate heterogeneity^14,15,30,31^. Consistent with this view, recent lineage-tracing studies in monoclonal gastruloids reveal that fate biases can emerge from heritable fluctuations in stem cell states even before induction, leading to divergent lineage outcomes in otherwise clonal aggregates^87^.

Single-cell transcriptomic analysis revealed a surprisingly mature posterior domain including neural crest, neural ectoderm, presomitic mesoderm, and somites. This finding indicates that, supplemented by temporally controlled retinoic acid exposure, this model could be adapted to continue development through neurulation^37,53,90^. Additionally, we see canonical markers of extraembryonic tissues, including Amnion (*ISL1*) and Yolk Sac (*APOA1*, *AFP*), suggesting that cross talk between embryonic and extraembryonic lineages can emerge autonomously^22,99^. Overall, our system recapitulates symmetry breaking, spatial segregation of the three germ layers, and complex cell types matching that of CS7 embryos^44^.

A key mechanistic finding is that WNT-driven heterogeneity generates divergent TGFβ/ACTIVIN/NODAL signaling dynamics, which in turn instruct the divergent formation of the mesoderm and endoderm. Functional perturbations indicated that sustained TGFβ signaling favors endoderm specification, whereas transient signaling promotes mesoderm^66,100^. This temporal logic is supported by a broad convergence of recent works, wherein WNT establishes mesendoderm competence, after which duration of NODAL/ACTIVIN and WNT activity act antagonistically to resolve the mesoderm or endoderm branchpoint^3,61,101,102^. This principle was confirmed across micropatterned colonies^103,104^, and 3D gastruloids^105,106^. To complement elegant work based on addition of diffusible agonists to aggregates^3,31,104^, in this optogenetic system ACTIVIN/NODAL dynamics emerged endogenously through WNT driven autocrine and paracrine signaling within the aggregate, demonstrating that heterogeneous WNT activation alone was sufficient to initiate the full signaling cascade. Single-cell RNA sequencing revealed a striking complementarity where definitive endoderm is enriched for *SMAD2* and TGFβR1, whereas mesoderm showed an increased expression of TGFβ ligands (TGFB1/2) and *SMAD1/5,* a divergence shaped by competition between type I receptors for shared type II receptors^107,108^.

This complementary receptor-ligand architecture suggests an emergent paracrine signaling axis in which mesoderm-derived TGFβ ligands reinforce SMAD2-dominant signaling competence in adjacent endodermal cells. Consistent with this model, primitive-streak like cells retain ACTIVIN type 2 receptors (ACVR2A/B) while downregulating TGFβ type 2 receptors (TGFβR2/3), creating differential ligand responsiveness despite shared SMAD 2/3 downstream signaling^102^. This suggests that the TGFβ-TGFβR axis becomes selectively restricted to endoderm as mesoderm loses the capacity to respond to its own paracrine ligands.

A second organizing principle is an adhesion code linking cell fate and tissue mechanics. Cadherins are essential in embryonic and neural morphogenesis and drive autonomous cell sorting in embryos and gastruloids with combinatorial patterns shaping sorting dynamics and axis formation^78,82,109–112^. In mouse gastruloids the E-to N-cadherin switch is critical for organization, but N-cadherin itself is dispensable, with its loss producing trunk like structures rather than abolishing patterning^83^. Our optoWnt gastruloids show the importance of N-cadherin, as its knockdown in optoWnt cells impaired mesendodermal segregation, replacing hemispherical organization with radial segregation. These findings are supported by zebrafish gastrulation, where N-cadherin is necessary for endoderm internalization and coordinated inward migration^113^. Beyond their structural role in cell sorting, cadherins may also influence fate decisions through their regulation of epithelial integrity. Epithelial disruption has been shown to be a prerequisite for TGFβ protein sensing in human pluripotent stem cells, raising the possibility that the WNT-driven cadherin switch in our system generates a competence gradient where variation in the timing of epithelial disruption creates heterogeneous TGFβ signaling duration, contributing to the mesoderm-endoderm bifurcation^114^. Other cadherins displayed lineage-biased expression, including *CDH3* enriched in mesoderm and *CDH11* enriched in both mesodermal and endodermal lineages^83^. These studies establish germ layer organization in human gastruloids is dependent on cadherin expression, in which N-cadherin is a critical driver of mesendodermal segregation, and suggest that a full adhesion code may involve combinatorial interactions of multiple cadherins.

In conclusion, our activation approach, which emulates stochastic WNT activation in a subpopulation of pluripotent cells, demonstrates the utility of optogenetic systems to understand and engineer symmetry breaking and morphogenesis in hPSC gastruloid models. By capturing key hallmarks of embryonic development, we offer a reproducible, scalable tool to understand and perturb molecular mechanisms of development. Future work may explore how selective WNT sensitivity and/or activation may be present and distributed preceding human gastrulation. Prevailing hypotheses include differential response to morphogen signaling^55,115^, maternal β-catenin^116^, and/or the biophysical role of embryo morphogenesis^117,118^, though these possibilities have yet to be determined. Furthermore, addition of extraembryonic lineages including Extraembryonic Endoderm (XEN) and Trophoblast (TS) stem cells would create a more representative model to interrogate additional complex signaling networks^78^. Additionally, light control of other major morphogen systems has recently been reported, including optoFGF,^119^ optoBMP^120^, optoNodal^70^, optoTGFβ^121^, and optoE-Cad^122^ which may enable future studies into the spatiotemporal role of morphogen signaling in hPSC models of development. Overall, these results motivate the use of selective WNT co-culture systems as an approach to investigate the mechanisms and morphogenic events underlying human gastrulation.

## Acknowledgments

We thank Todd McDevitt, Kyle Loh, Richard Harland, and Valerie Weaver for helpful discussions on stem cell and developmental biology. We also thank members of the David Schaffer lab for helpful discussions throughout the project. We are grateful to Mary West from the QB3 High-Throughput Screening Facility and CIRM/QB3 Shared Stem Cell Facility and Justin (Yoo Gi) Choi from QB3 Genomics for technical assistance.

## Author contributions

Conceptualization: H.J.J., J.A.Z., T.N., and D.V.S.

Methodology: H.J.J., J.A.Z., N.A.R., and D.V.S.

Investigation: H.J.J., D.M.M., J.A.Z., C.N.K., N.A.R., R.B., and D.V.S

Visualization: H.J.J., D.M.M., J.A.Z., C.N.K., N.A.R., R.B., and D.V.S

Funding acquisition: D.V.S. Project administration: D.V.S. Supervision: H.J.J., and D.V.S.

Writing – original draft: H.J.J, J.A.Z., and D.V.S.

Writing – review & editing: H.J.J, D.M.M., T.N., and D.V.S.

## Inclusion and diversity statement

*will add if accepted in principal*

## Methods

### Generation of hESC optoWnt cell lines

The hESC optoWnt cell lines were generated as previously described by Repina and Johnson^40^.

### hESC cell culture

For routine culture and maintenance, all optogenetic and WT hESC lines (H9, WiCell) and were grown on Matrigel (Corning, lot # 7268012, 7275006) coated plates in mTeSR1 medium (STEMCELL Technologies) and 1% penicillin/streptomycin (Life Technologies) at 37 °C and 5% CO2 with daily media changes. Optogenetic cells were cultured with hood lights off. For 2D well plate illumination experiments, cells were singularized with Accutase (STEMCELL Technologies) at 37 °C for 5 min and seeded onto Matrigel-coated 96-well plates in media containing 10 µM ROCK inhibitor Y-27632 (Selleckchem). Cells were seeded at a density of 35k cell cm-2. For co-culture experiments, WT and optoWnt cells were mixed in a 1:1 ratio and seeded at a density of 35k cell cm-2. After 20-24 hrs, media was changed to either mTeSR1 or growth factor reduced medium (GFM) without ROCK inhibitor and plates were placed onto LAVA illumination devices and subjected to experimental conditions. Growth factor reduced (GFR) medium consisted of DMEM-F12 (STEMCELL Technologies) with N2 (1:200, Invitrogen) and B27 without vitamin A (1:100, Invitrogen).

For 3D cellular aggregate preparation, cells were singularized with Accutase (STEMCELL Technologies) at 37 °C for 5 min and seeded onto AggreWell 400 plates at a density to form 200 cells/aggregate, centrifuged at 200g for 4 min at room temperature, and compacted overnight at 37 °C and 5% CO2 in mTeSR1 medium (STEMCELL Technologies) plus 10 µM ROCK inhibitor Y-27632 (Selleckchem). After 20-24 hours of compaction, the cell aggregates were collected, washed, and seeded in a thermoreversible PEG-PNIPAAm hydrogel (Mebiol, CosmoBio) in the liquid phase at 4 °C at 10 wt%. Gel droplets of 50 uL were formed containing ∼200 well dispersed 3D cell aggregates in 24 well glass bottom plates (Eppendorf) and heated at 37 °C for 10 min to form transparent hydrogels. Prewarmed GFR media was then added to each well, the plate was placed onto the LAVA illumination devices and subjected to experimental conditions. To collect the aggregates for downstream analysis, cold PBS was added to each well and the plate was incubated at 4C for 10 minutes, to liquify the thermoreversible hydrogels and release the 3D cell aggregates.

### Cadherin Knockdowns

To generate shRNA knockdown lines, shRNA sequences (Table S1) were subcloned into the pLKO.1 lentiviral expression vector digested with AgeI and EcoRI, and modified to express the blastocydin-resistance gene and eGFP. Knockdown lines were generated by lentiviral infection of optoWnt hESCs with shRNA against target genes. Infected cells were isolated by FACS sorting for eGFP expression. Knockdown was verified through western blot for target genes.

### Optogenetic stimulation

Cells plates were placed onto engineered illumination devices, described previously^45^ and maintained in standard 37°C tissue culture incubators. In brief, user-defined illumination patterns were uploaded to the engineered illumination device for independent illumination control of each well. Unless otherwise noted, optogenetic stimulation was achieved with blue light emitted by arrays of 470nm LEDs continuously illuminating hESCs with 0.8 µW mm-2 light for the duration of the experiment (1-48 hrs).

### Immunostaining and imaging

For 2D and 3D cell cultures, cells were fixed with 4% paraformaldehyde (ThermoFisher) in PBS for 20 min (2D) or 30 min (3D) at room temperature and subsequently washed three times with PBS, followed by blocking and permeabilization with 5% donkey serum (Sigma-Aldrich) and 0.3% Triton X-100 (Fisher Scientific) in PBS (PBS-DT) for 1 hour. Cells were incubated with primary antibodies at 4 °C overnight, then washed three times with PBS, and incubated with fluorescently conjugated secondary antibodies (Invitrogen) at 1:250 dilution for 1 hour at room temperature. Both primary and secondary antibodies were diluted in PBS-DT. Cells were washed with PBS and stained with 0.1 µg mL −1 DAPI nuclear stain (ThermoFisher) prior to imaging. Confocal imaging was performed on a Perkin Elmer Opera Phenix system (QB3 High-Throughput Screening Facility). Brightfield and widefield fluorescence imaging was performed on a Molecular Devices Image Xpress Micro imaging system (CIRM/QB3 Shared Stem Cell Facility).

### Image analysis

Line trace analysis and aggregate reproducibility plots were generated from a custom Matlab script using the .tiff files of the maximum intensity projections for each aggregate, collected at 20X from the Opera Phenix system. Generally, the DAPI stain was used to classify each cell nuclei, and assigned as marker positive for either *SOX2*, *BRA/T*, or *SOX17* or marker negative, based on fluorescence intensity. The spatial distribution of teach classified nuclei was then projected to a single vector axis, drawn from the greatest separation between the two populations, and the resulting line trace analysis was plotted based on the normalized diameter for each aggregate. Lines represented average % of cells maker positive along the distance of the axis, pooled from multiple aggregates. Error bars represented the standard deviation of the aggregate population.

### RNA extraction, reverse transcription, and qPCR

Cells were lifted with Accutase at 37 °C for 5min, centrifuged, and resuspended in TRI reagent (Zymo Research). To achieve higher RNA yields, two to three wells were pooled, constituting a single biological replicate. RNA was purified using an RNA extraction kit (Zymo Research) as per manufacturer recommendations with an on-column DNase digestion to remove residual genomic DNA. After measurement of total RNA concentration, 1µg of RNA was converted to cDNA using an iScript cDNA synthesis kit (Bio-Rad). Finally, 10ng of cDNA was used for each SYBR Green qPCR reaction, run in 96-well plate format with a 0.1µM final forward and reverse primer concentration. qPCR was conducted for 40 cycles at an annealing temperature of 56 °C on a CFX Connect Real-Time PCR Detection System (Bio-Rad). A melt curve was generated at the end of the PCR reaction and a subset of reactions were run on a 1% agarose gel to ensure that only one product of the expected size was amplified per primer pair. qPCR analysis was conducted by the ddCt method. For each cDNA sample, gene expression was internally normalized to the expression of a human housekeeping gene (GAPDH or ACTB) run on the same qPCR plate. Next, for each gene, expression was normalized to the expression level of WT untreated hESCs. The log of relative expression over this WT control (i.e. log2(fold change)) was graphed as a heatmap where color corresponds to mean value of biological replicates. The variability in gene expression was assessed with histogram graphs that show mean and standard deviation of fold change for 3 biological replicates, with at least 2 technical replicates for each biological replicate.

### Single cell RNA sequencing and data analysis

Library preparation, sequencing, and data pre-processing were performed by QB3 Genomics at the University of California, Berkeley. In short, 10x genomics libraries were sequenced using NovaSeq 6000 (Illumina), and FASTQ files were assessed for sequence quality with FastQC (v.0.12.0). CellRanger was used to create an initial cell by gene matrix. STARsolo v2.7.11^123^ was used to align reads to the Human Genome GRCh38. This matrix was converted to an anndata object and processed using scanpy^124^. Solo was used for doublet detection and removal with default parameters and a 0.5 cutoff for the softmax/doublet score^125^. Quality control steps included filtering low quality cells based on a minimum of 2000 unique genes, a 12% mitochondrial cutoff, and a 30% ribosome cutoff. Counts were normalized to 10,000 counts per cell, log transformed and regressed to remove contribution of ribosomal and mitochondrial related genes. Dimensionality reduction was performed using Uniform manifold approximation and projection (UMAP) embeddings^126^ calculated with the top 30 principal components. Initial clustering was performed with Leiden clustering^127^ and followed by final clustering with k-means. To validate cluster annotations we integrated CS7 Human Gastrula^44^ using QC cutoff of 2000 unique genes, a 12% mitochondrial cutoff. To integrate the two datasets sciVI^128^ was used to reduce batch effects. Additionally, pseudocounts of each cell type cluster from the optoWNT dataset was averaged creating a single pseudo-cell that was then integrated into the analysis.

For RNA velocity analysis spliced and unspliced count matrix were also produced with STARSolo and used for quantification. scVelo was used for RNA velocity with the dimensional reduction for PCA and UMAP using the spliced count matrix only and cluster annotations. The dynamical model was used in the first pass and refitted on the top 100 genes computed from differential kinetics testing on the cluster annotations. Latent time representation of the RNA velocity model was used to order the cells for visualization of gene expression signatures. Dpt pseudotime was run by selecting a cell at random from the epiblast cluster and pseudotime values were assigned to cells relative to this root. Cell signaling through ligand-receptor analysis was performed using CellChat(v2.1.2)^72^. In R we converted the anndata object with cell type specific labels to a CellChat to perform further downstream analysis.

### Statistical analysis and graphing

Data are presented as mean ± 1 standard deviation (s.d.) unless otherwise specified. Statistical significance was determined by Student’s t-test (two-tail) between two groups, and three or more groups were analyzed by one-way analysis of variance (ANOVA) followed by Tukey test. The unpaired two-samples Wilcoxon test was performed for data that was not normally distributed. P < 0.05 was considered statistically significant (NS P>0.05, *P<0.05, **P<0.01, ***P<0.001). Statistical analysis and data plotting was performed in R.

### Data Availability

All raw data used here are publicly available. For alignment of sequencing data GRCh38() was used. The sequencing data that supports the findings of this study have been deposited in the Gene Expression Omnibus. Previously published sequencing data that were re-analyzed are available under the following ascension code. All other data supporting the findings of this study are available upon request. A github repository will be cited to reproduce the analyses.

