## Supplementary Information for "Optogenetic WNT signaling drives germ layer self-organization in a human gastruloid model"

**This PDF file includes:**

Supplementary Text Figs. S1 to S12
Supplementary Table 1

### Supplementary Figures

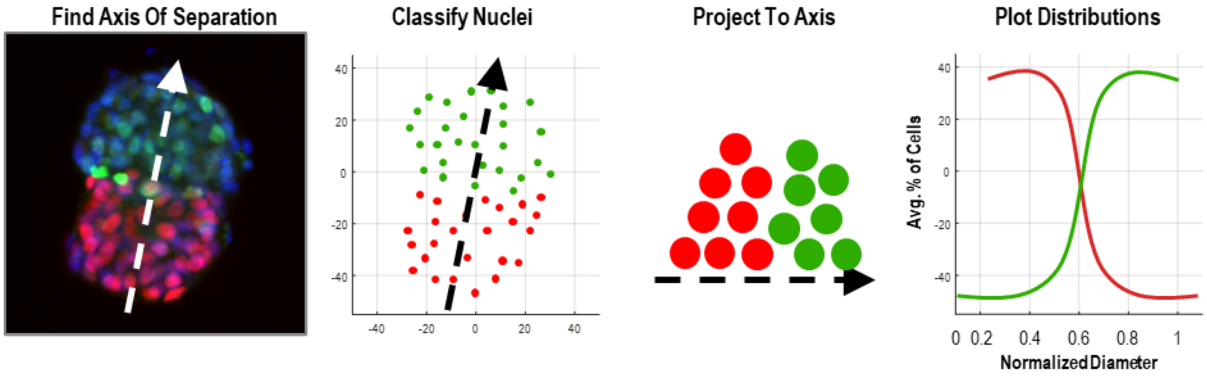


**Fig. S1.**

**Schematic of cellular line-trace analysis**

Image analysis performed in Matlab custom script (see methods)

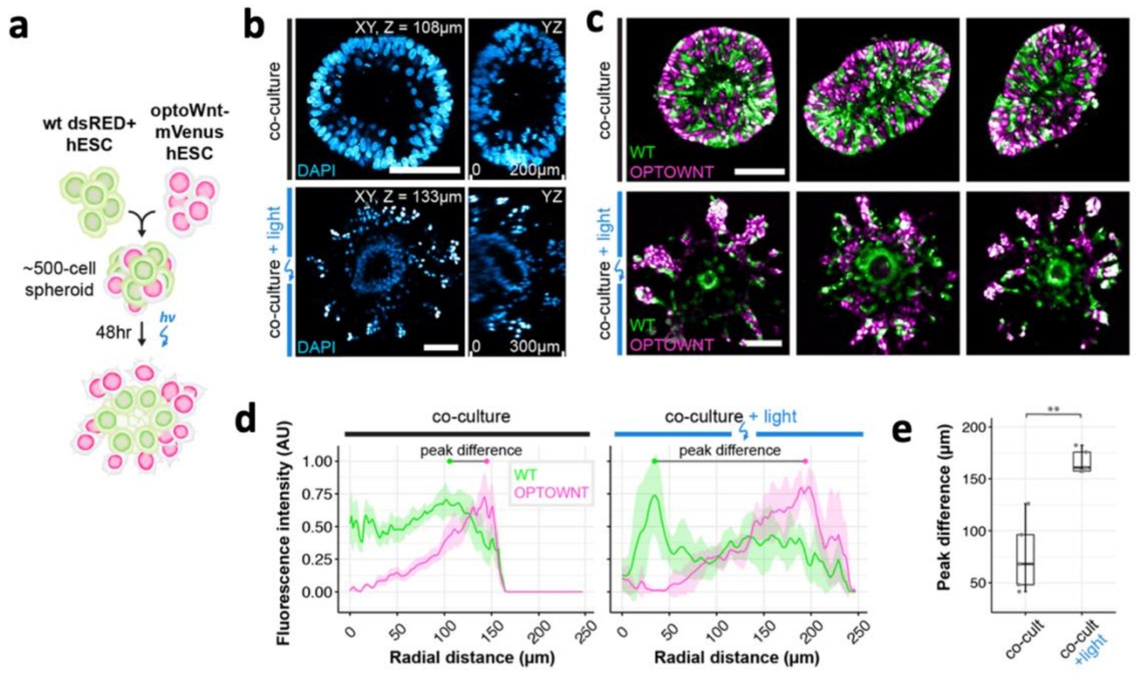


**Fig. S2. optoWnt-induced cell organization of hESC aggregates in Matrigel**

**a,** Schematic of experimental setup of optoWnt/WT co-cultures in 3D spheroid cultures. **b,** Two photon imaging of co-cultures kept in the dark (top) and after 48 hrs illumination (bottom) with DAPI staining of cell nuclei. Scale bars 100 µm, YZ axial cross-sections 200 µm in height. **c,** Two-photon imaging of co-cultures kept in the dark (top) and after 48 hrs illumination (bottom). WT cells are labelled with dsRed, optoWnt cells are labelled with mVenus-NLS expression. Scale bars 100µm. **d,** Radial quantification of cell segregation in 3D spheroids. Normalized intensity of mean fluorescence signal is graphed as a function of radial distance from spheroid center. Graph shows mean of n = 5 spheroids, error bars = 1 S.E.M. E) Radial distance between peak intensities of WT and optoWnt fluorescence distribution, i.e. 'peak difference', labelled in **e,** Unpaired two-samples Wilcoxon test (p = 0.0079), n = 5 spheroids.

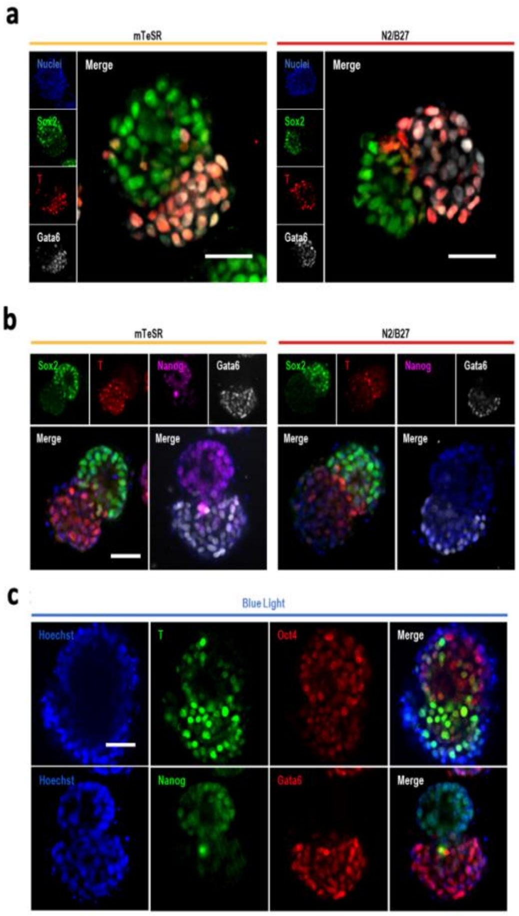
a
mTeSR
N2/B27
Nuclei
Merge
Sox2
T
Gata6
b
Nanog
c
Blue Light
Hoechst
Oct4

**Fig. S3. Expression of pluripotency and endoderm markers in the 60-hour optoWnt gastruloids. a,** Representative maximum intensity projection images of 60-hour optoWnt gastruloids immunostaining for markers of the ectoderm (Sox2), mesoderm (T), and endoderm (Gata6) in either mTeSR pluripotency media or basal media containing N2/B27 supplement. Mesendoderm specification observed in both media types. **b,** Representative images of 60-hour optoWnt gastruloids immunostaining for markers of the ectoderm (Sox2), mesoderm (T), pluripotency (Nanog), and endoderm (Gata6) in either pluripotency media (mTeSR) or basal media (N2/B27). Robust segregation occurs in both medias, with loss of Nanog expression in the GFR media. Scale bar = 50 µm. **c,** Representative maximum intensity projection images of 60-hour gastruloids immunostaining for markers of the mesoderm (T), pluripotency (Oct4, Nanog), or endoderm (Gata6). Pluripotency markers and mesoderm/endoderm markers are expressed in different hemispheres of the segregated gastruloids. Scale bar = 50 µm.

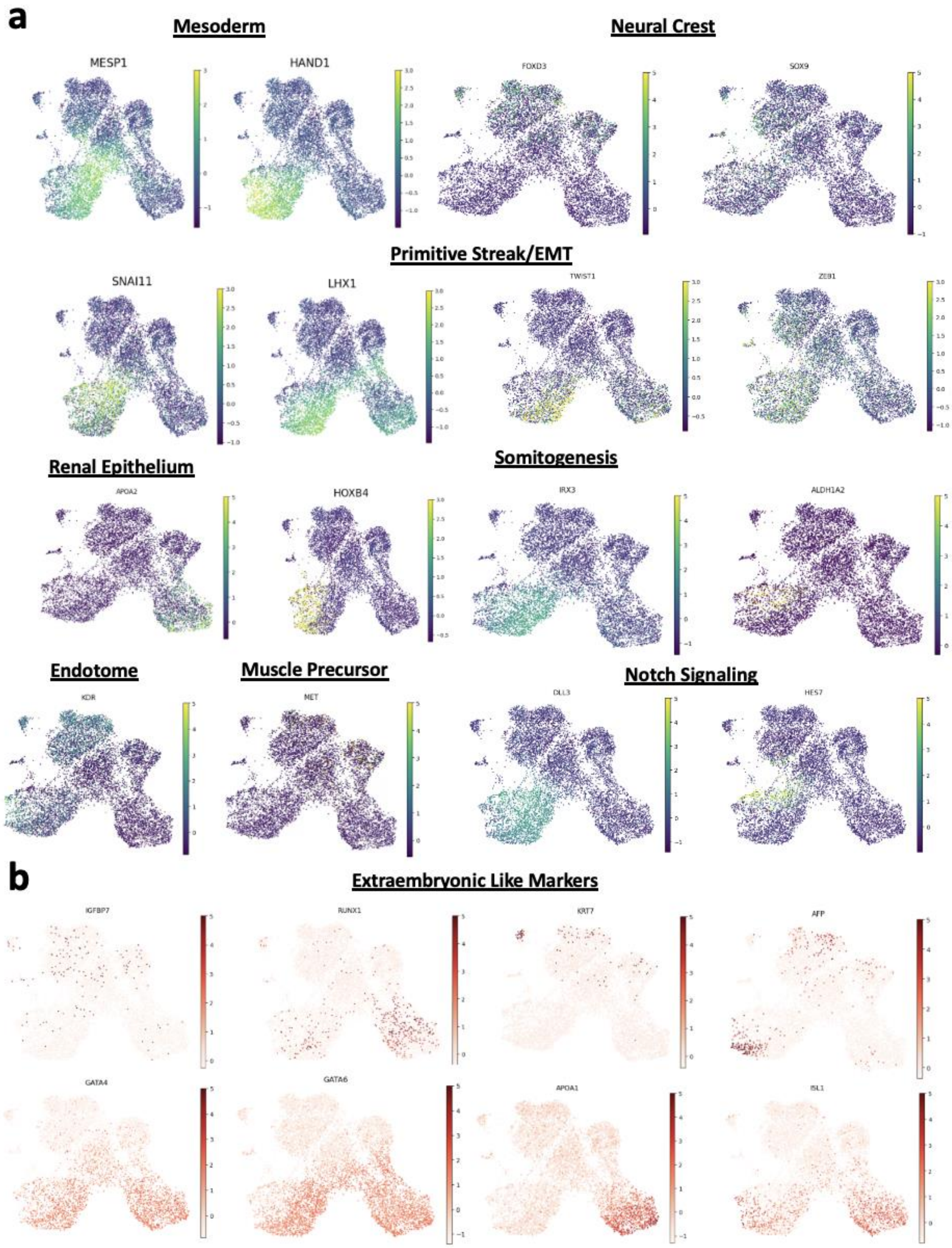


**Fig. S4. Expression of pluripotency markers in the 60-hour optoWnt gastruloids. Fig. S4. Evaluation of cell type expression profiles in 60 hour optoWNT gastruloids.** a, UMAP projections of normalized expression of lineage-specific markers across identified cell populations in 60-hour optoWNT sc-RNAseq data. Markers shown include mesoderm (MESP1, HAND1), neural crest (FOXD3, SOX9), primitive streak/EMT (SNAI11, LHX1, TWIST1, ZEB1), renal epithelium (APOA2, HOXB4), somitogenesis (IRX3, ALDH1A2), endotome (KDR), muscle precursor (MET), and Notch signaling (DLL3, HES7). b, Normalized expression of canonical extraembryonic lineage markers (IGFBP7, RUNX1, KRT7, AFP, GATA4, GATA6, APOA1, ISL1) on UMAP projections.

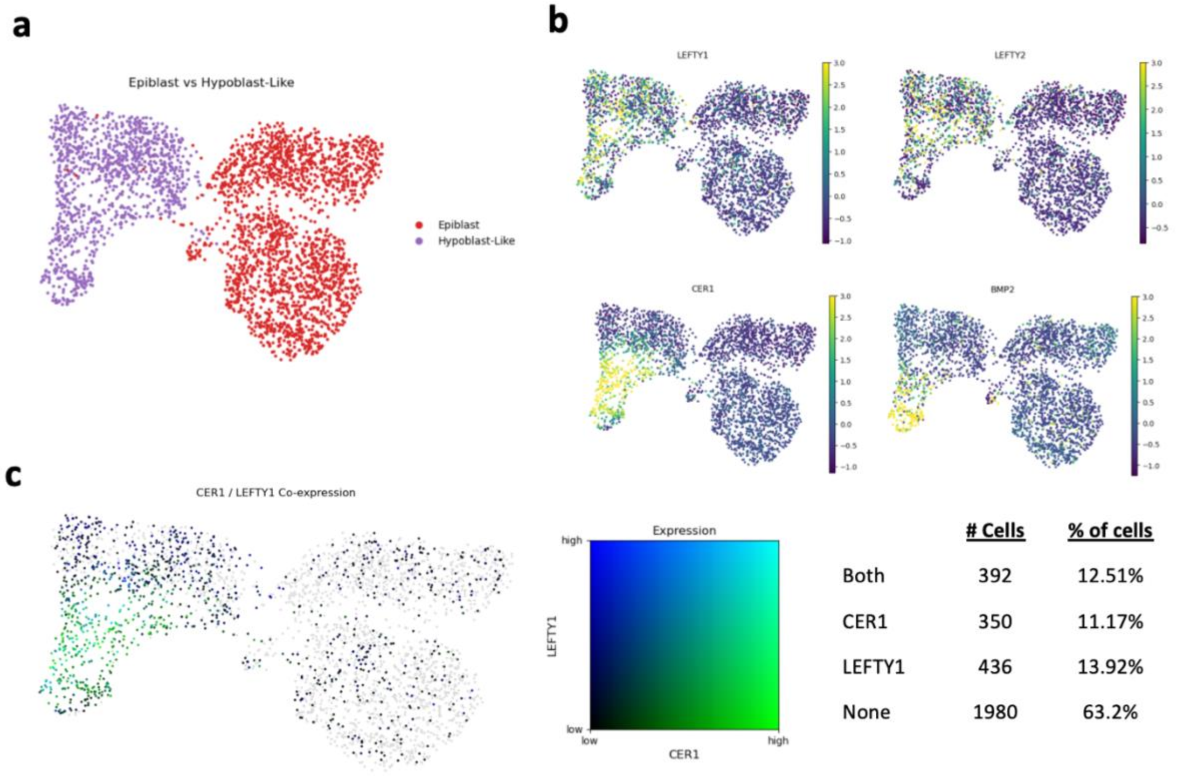


**Fig. S5 Characterization of epiblast and hypoblast-like populations in 60-hour optoWNT gastruloids. a,** UMAP projection of cells annotated as epiblast and hypoblast-like, subset from the full dataset using cell type annotations defined in Fig. 2b. **b,** UMAP projections showing normalized expression of hypoblast-associated markers (LEFTY1, LEFTY2, CER1, BMP2) across the epiblast and hypoblast-like populations. **c,** Co-expression analysis of CER1 and LEFTY1. UMAP colored by joint CER1/LEFTY1 expression on a per-cell basis. (Left) two-dimensional colormap indicating co-expression intensity. (Middle) quantification of cells positive for both markers (12.51%), CER1 only (11.17%), LEFTY1 only (13.92%), or neither (63.2%). (Right)

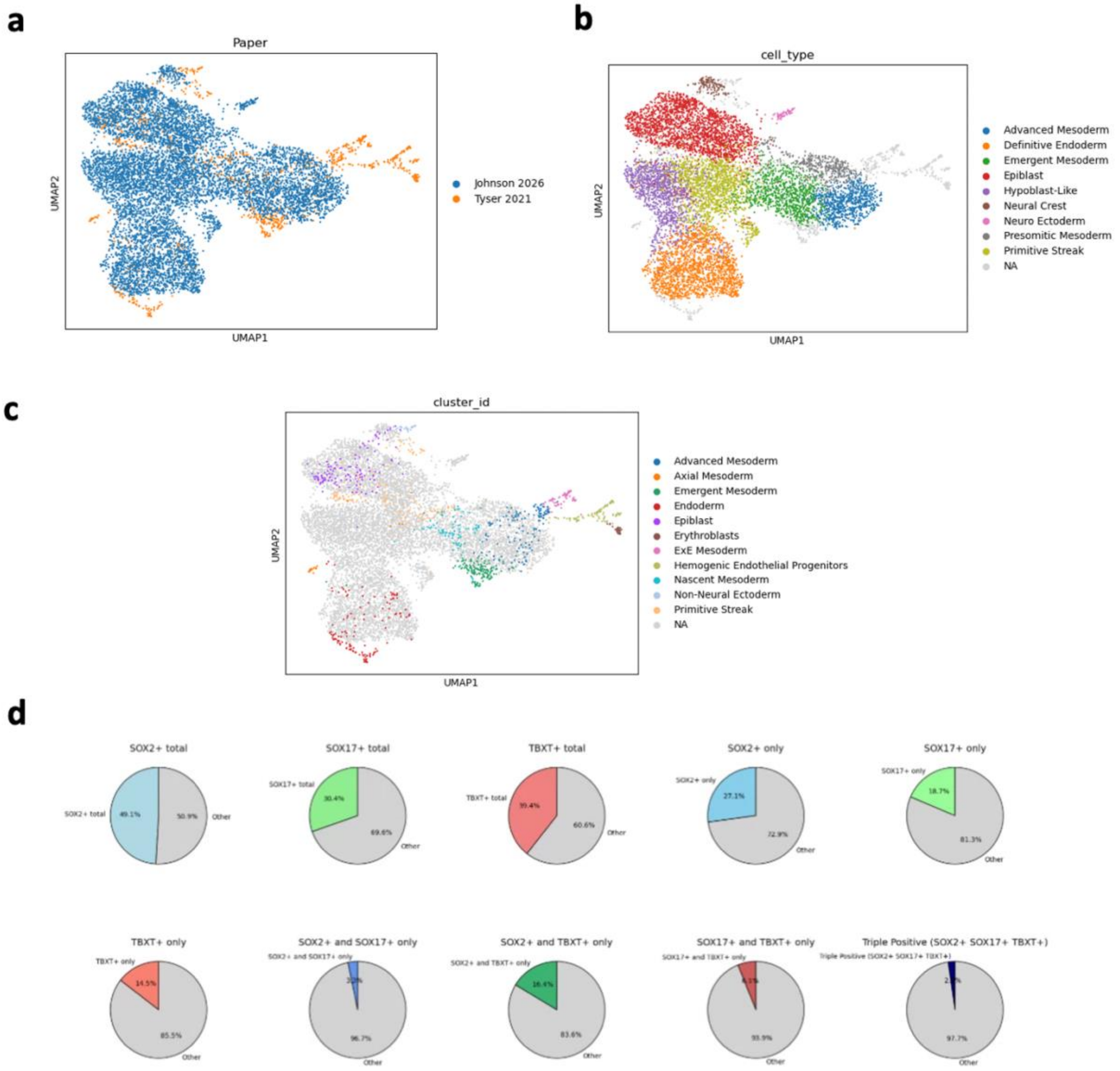


**Fig. S6.**

**Co-embedding of optoWNT gastruloid sc-RNAseq profiles with published data from human embryos a,** UMAP projection of integrated sc-RNAseq data from 60-hour optoWNT gastruloids from this study (Blue) and Tyser 2021 CS7 human embryo (orange). **b,** Co-embedded UMAP colored by cell type annotations from the 60-hour optoWNT gastruloid dataset. **c,** Co-embedded UMAP colored by cell type annotations from the CS7 human embryo dataset. d, Pie charts displaying the proportion of cells expressing SOX2, SOX17, and TBXT, either individually or in combination, from all cells in the 60-hour optoWNT sc-RNAseq data..

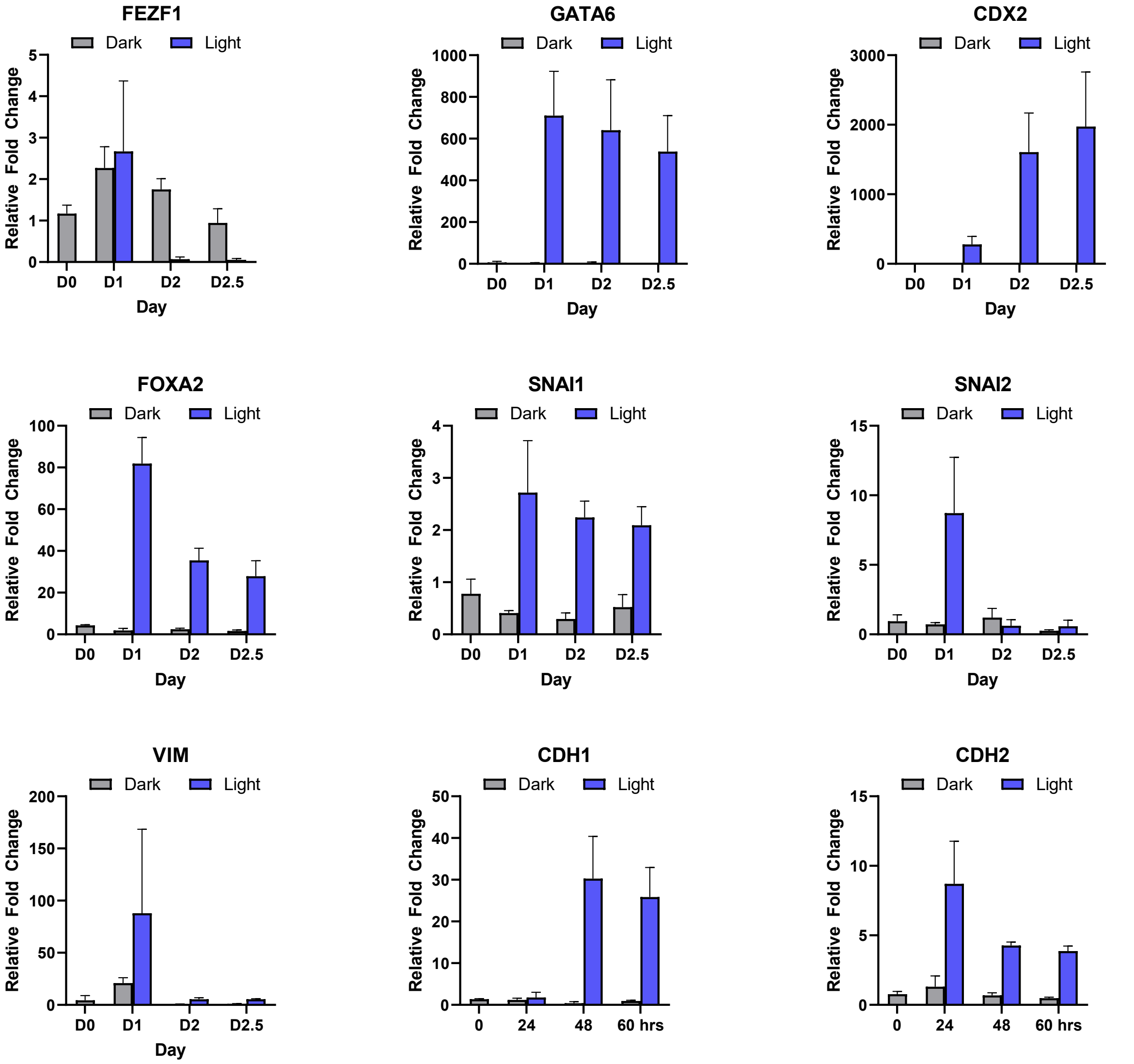


**Fig. S7. optoWnt Gastruloid gene expression vs time as measured by qPCR.** qPCR for markers of the 3 germ layers and EMT in optoWnt gastruloids from hour 0 to hour 60. Graphs show relative fold change as compared to hESCs. n=3 biological replicates of pooled gastruloids. Error bars = std. dev.

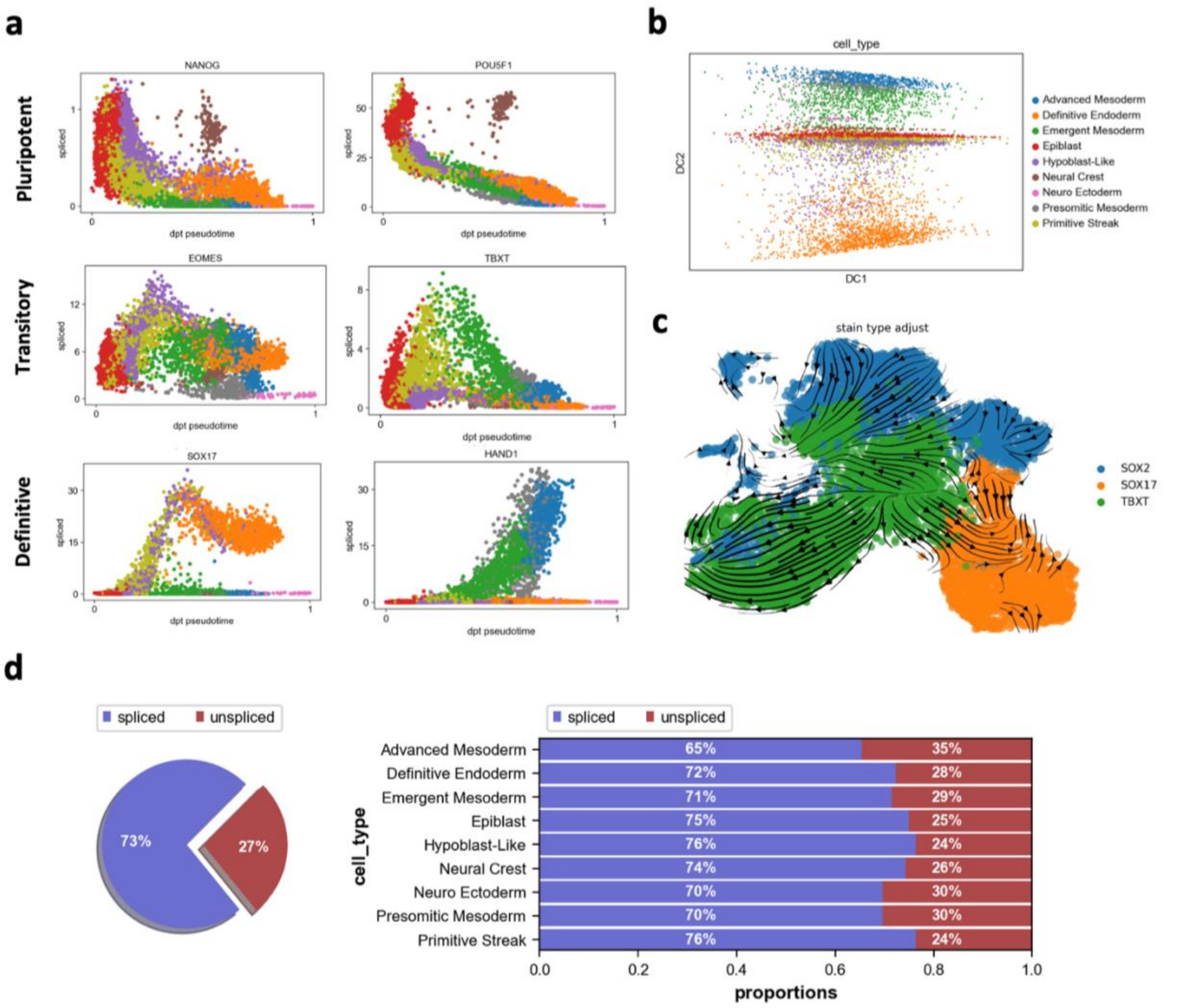


**Figure S8. Additional RNA velocity analysis of 60-hour optoWnt gastruloids with continuous blue light illumination. a,** Spliced transcript expression plotted against diffusion pseudotime (dpt) for representative genes across three categories: pluripotent (NANOG, POU5F1), transitory (EOMES, TBXT), and definitive (SOX17, HAND1) **b,** Diffusion map projection (DC1 vs DC2) colored by cell type. **c,** RNA velocity vectors overlaid on a UMAP projection, with cells colored by the highest expressing germ layer marker (SOX2, ectoderm; SOX17, endoderm; TBXT, mesoderm), **d,** Proportion of spliced versus unspliced transcripts globally (left, pie chart) and per cell type (right, stacked bar chart).

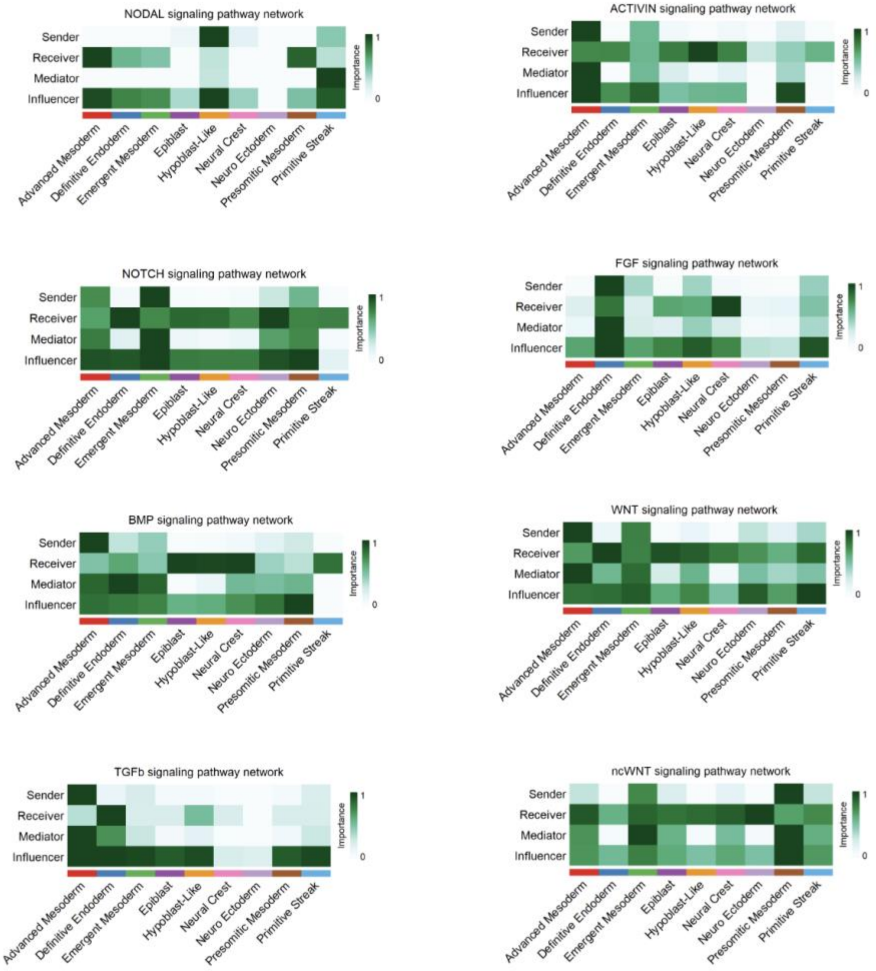


**Fig. S9**. **CellChat signaling network analysis of 60-hour optoWNT gastruloids**. Heatmaps displaying inferred cell-cell communication roles across identified cell type clusters for major developmental signaling pathways. Color intensity represents a relative importance score (0–1) for each cell type within each signaling role, as computed by CellChat. See Methods.

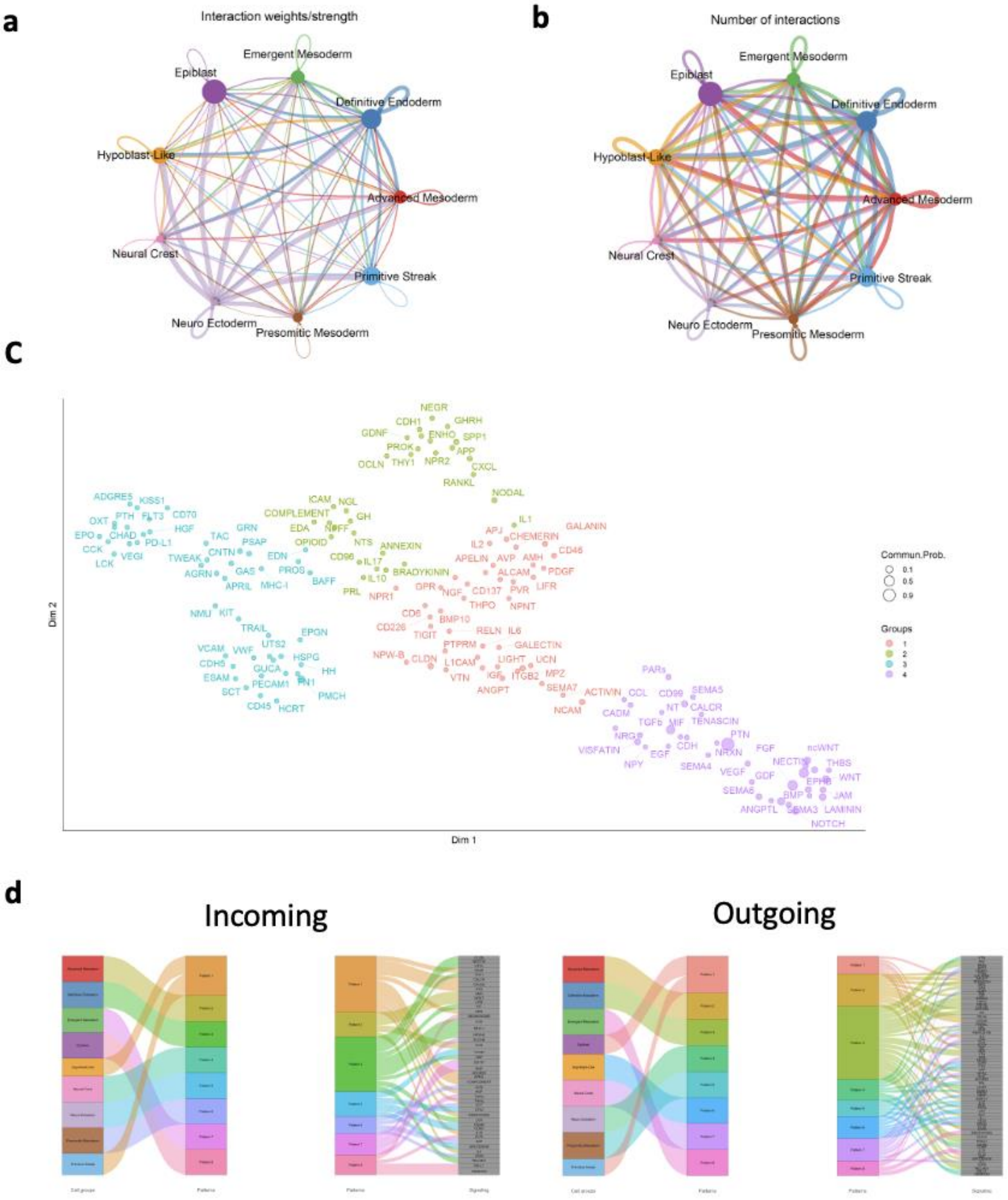


**Fig. S10. Cellchat overall network analysis a,** Chord plot for total number of inferred interactions for each cell type **b,** Chord plot for weighted aggregate cell-cell communication network showing the interaction strength **c,** Manifold learning analysis of signaling network clustering based on functional similarities **d,** River plots of inferred signaling patterns mapped to specific cell groups (Left) and (Right) for both incoming and outgoing signaling

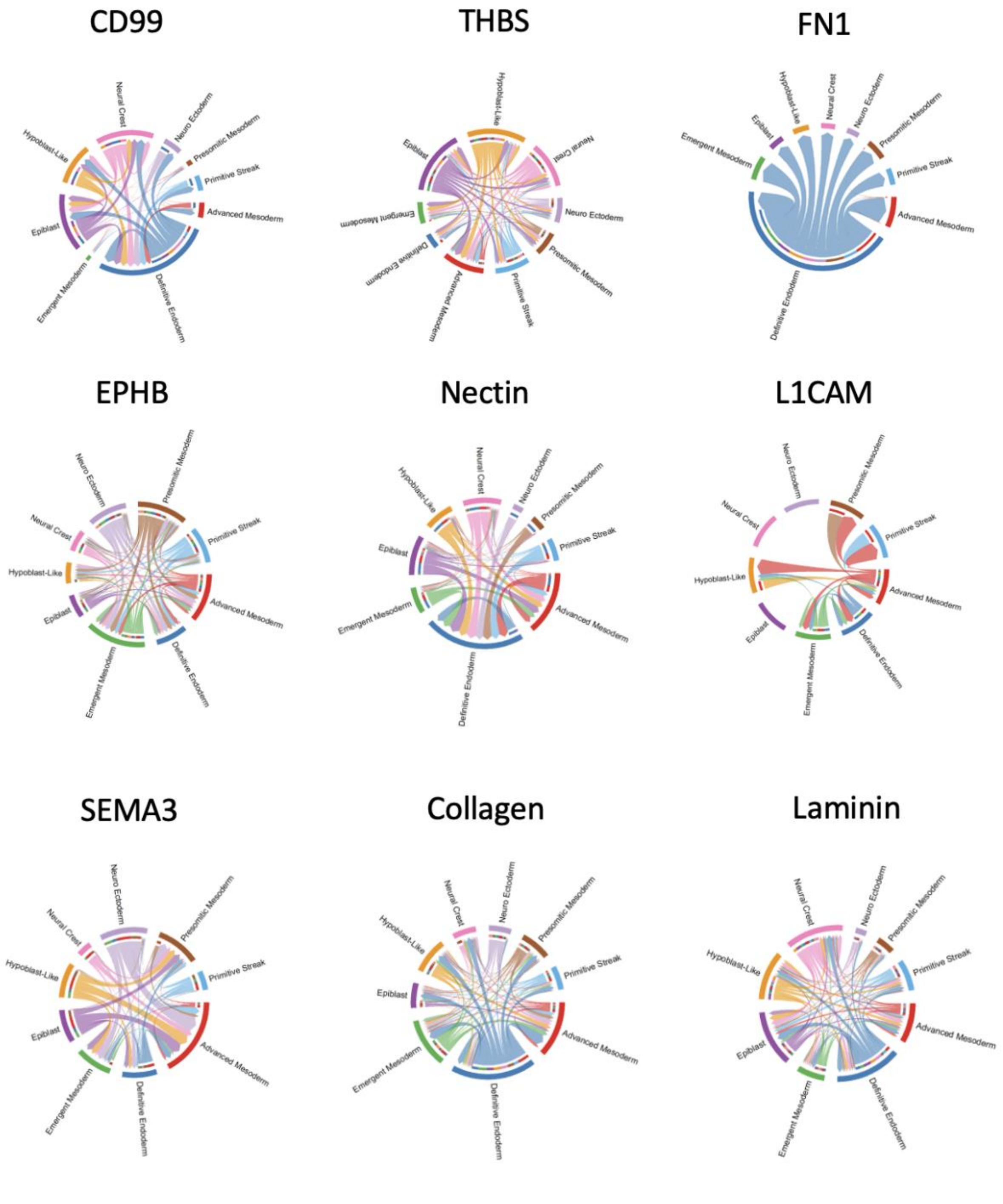


**Fig. S11. Cellchat Cell Analysis of Adhesion Molecules and Extracellular Matrix** Chord diagrams depicting inferred cell-cell communication networks.

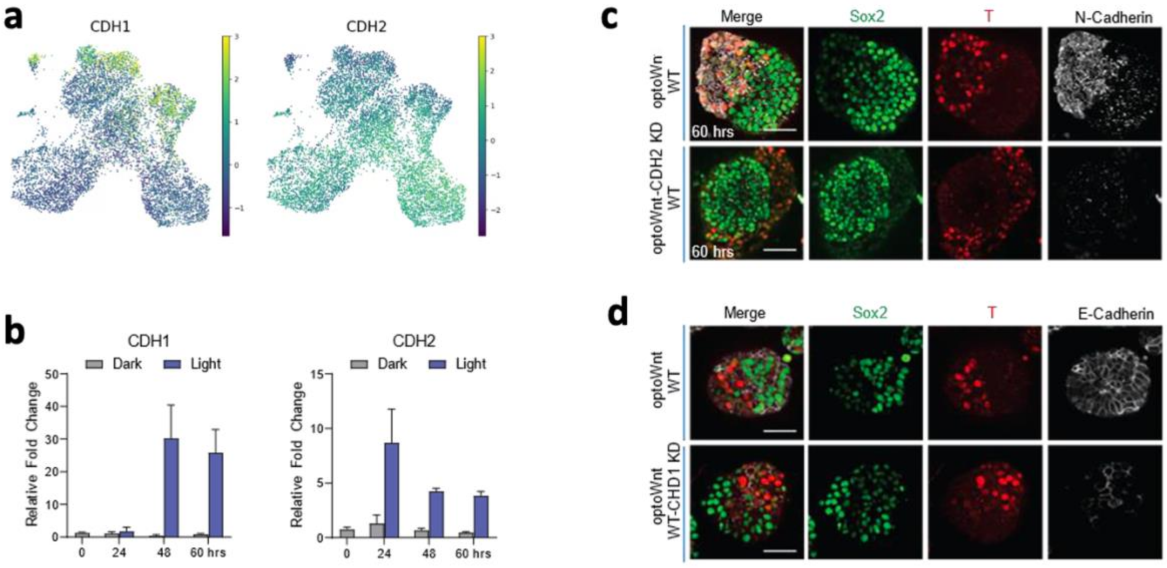


**Fig. S12. E-Cadherin and N-Cadherin gene expression and knock down generation in optoWnt gastruloids. a,** Single cell RNA sequencing analysis of 60-hour optoWnt gastruloids with relative expression of E-Cadherin (CDH1) and N-Cadherin (CDH2) within the UMAP clustering projections. Sample represents a pooled population of ~50 optoWnt gastruloids. **b,** qPCR of CDH1 and CDH2 at various time points of optoWnt gastruloid patterning. Graph represents relative fold change with respect to hESCs, n=3 biological replicates of pooled gastruloids. Error bars = s.d. **c,** Representative maximum intensity projection images of 60-hour optoWnt gastruloid N-Cadherin expression and loss of expression in the optoWnt-Ncad KD line. **d,** Representative images of 60-hour optoWnt gastruloid E-Cadherin expression and loss of expression in the optoWnt-Ecad KD line. Scale bar = 50 µm.
